# Expansion of a bacterial operon during cancer treatment ameliorates drug toxicity

**DOI:** 10.1101/2024.06.04.597471

**Authors:** Kai R. Trepka, Wesley A. Kidder, Than S. Kyaw, Taylor Halsey, Christine A. Olson, Edwin F. Ortega, Cecilia Noecker, Vaibhav Upadhyay, Dalila Stanfield, Paige Steiding, Benjamin G. H. Guthrie, Peter Spanogiannopoulos, Darren Dumlao, Jessie A. Turnbaugh, Matthew D. Stachler, Erin L. Van Blarigan, Alan P. Venook, Chloe E. Atreya, Peter J. Turnbaugh

**Author notes:** These authors contributed equally to this work (Kai R. Trepka, Wesley A. Kidder).

## Abstract

Dose-limiting toxicities remain a major barrier to drug development and therapy, revealing the limited predictive power of human genetics. Herein, we demonstrate the utility of a more comprehensive approach to studying drug toxicity through longitudinal study of the human gut microbiome during colorectal cancer (CRC) treatment (NCT04054908) coupled to cell culture and mouse experiments. 16S rRNA gene sequencing revealed significant shifts in gut microbial community structure during oral fluoropyrimidine treatment across multiple patient cohorts, in mouse small and large intestinal contents, and in patient-derived *ex vivo* communities. Metagenomic sequencing revealed marked shifts in pyrimidine-related gene abundance during oral fluoropyrimidine treatment, including enrichment of the *preTA* operon, which is sufficient for the inactivation of active metabolite 5-fluorouracil (5-FU). *preTA*^+^ bacteria depleted 5-FU in gut microbiota grown *ex vivo* and the mouse distal gut. Germ-free and antibiotic-treated mice experienced increased fluoropyrimidine toxicity, which was rescued by colonization with the mouse gut microbiota, *preTA*^+^ *E. coli*, or *preTA*-high CRC patient stool. Finally, *preTA* abundance was negatively associated with fluoropyrimidine toxicity in patients. Together, these data support a causal, clinically relevant interaction between a human gut bacterial operon and the dose-limiting side effects of cancer treatment. Our approach is generalizable to other drugs, including cancer immunotherapies, and provides valuable insights into host-microbiome interactions in the context of disease.

**One Sentence Summary:** Gut microbial enzymes can be used to predict and prevent anticancer drug toxicity.

## INTRODUCTION

Dose-limiting toxicities pose a major challenge in drug development (*1*). Decades of pharmacogenomic research have identified host alleles responsible for variable drug metabolism, yet clinical implementation has lagged due to persistent unexplained variability in toxicity (*2, 3*). The gut microbiome complements and extends host pathways of drug metabolism, with depletion of hundreds of pharmaceuticals following *in vitro* incubation with gut strains (*4, 5*). However, the impact of microbial drug metabolism on toxicity *in vivo* remains underexplored (*6*).

Despite advances in immunotherapy, cytotoxic chemotherapies such as fluoropyrimidines remain a cornerstone of gastrointestinal cancer treatment (*7*). A significant obstacle in fluoropyrimidine therapy is high interpatient variability in toxicity, necessitating dose adjustments in 35% of patients and therapy discontinuation in 10% (*8*). The widely-used oral fluoropyrimidine capecitabine (CAP) is a prodrug that is converted to 5-fluorouracil (5-FU), which kills cells through disruption of DNA synthesis and RNA processing (*9*). Toxic 5-FU is cleared to inactive dihydrofluorouracil (DHFU) by homologous host and microbial enzymes (DPYD and PreTA, respectively) (*10*). While rare patient DPYD sequence variants are associated with severe CAP toxicity (*11*), host genetic variation and other risk factors cannot explain extensive inter-subject variation in pharmacokinetics and toxicity (*12–14*). The gut microbiome varies widely between colorectal cancer (CRC) patients (*15*), offering a potential source of variability in toxicity. However, the impact of microbial *preTA* on predicting and preventing CAP toxicity remains unknown.

Here, we explore the role of microbial *preTA* in toxicity through the combined use of longitudinal studies of CRC patients receiving chemotherapy and follow-on studies *ex vivo* and *in vivo*. Together, our results highlight the importance of considering drug-microbe interactions to gain a more comprehensive view of mechanisms of variability in drug toxicity.

## RESULTS

### Oral fluoropyrimidines perturb gut microbial community structure

We conducted the <u>G</u>ut microbiome and <u>O</u>ral fluoropyrimidine (GO) clinical study (ClinicalTrials.gov ID NCT04054908), a prospective longitudinal study investigating the impact of oral fluoropyrimidines on the gut microbiome. Stool was collected at 7 timepoints spanning pre-treatment to post-cycle 3 of chemotherapy **(Fig. 1A)**. Of the 52 patients enrolled, 40 submitted at least one stool sample **(Fig. S1A)**. Patients were distributed among three sub-cohorts (**Table S1**): (A) CAP as monotherapy or part of a standard-of-care regimen (*n* = 21); (B) TAS-102 (trifluridine/tipiracil, *n* = 9); (C), CAP with bevacizumab and pembrolizumab immunotherapy (patients participating on ClinicalTrials.gov ID NCT03396926, *n* = 10). The 40 participants had a mean age of 52.5±10.5 y; 50% were male, 78% identified as white, and 8% identified as Hispanic or Latino ethnicity (**Table S1**). 83% of patients were TNM stage III-IV and 68% had prior surgery (**Table S1**).

**Figure 1:**
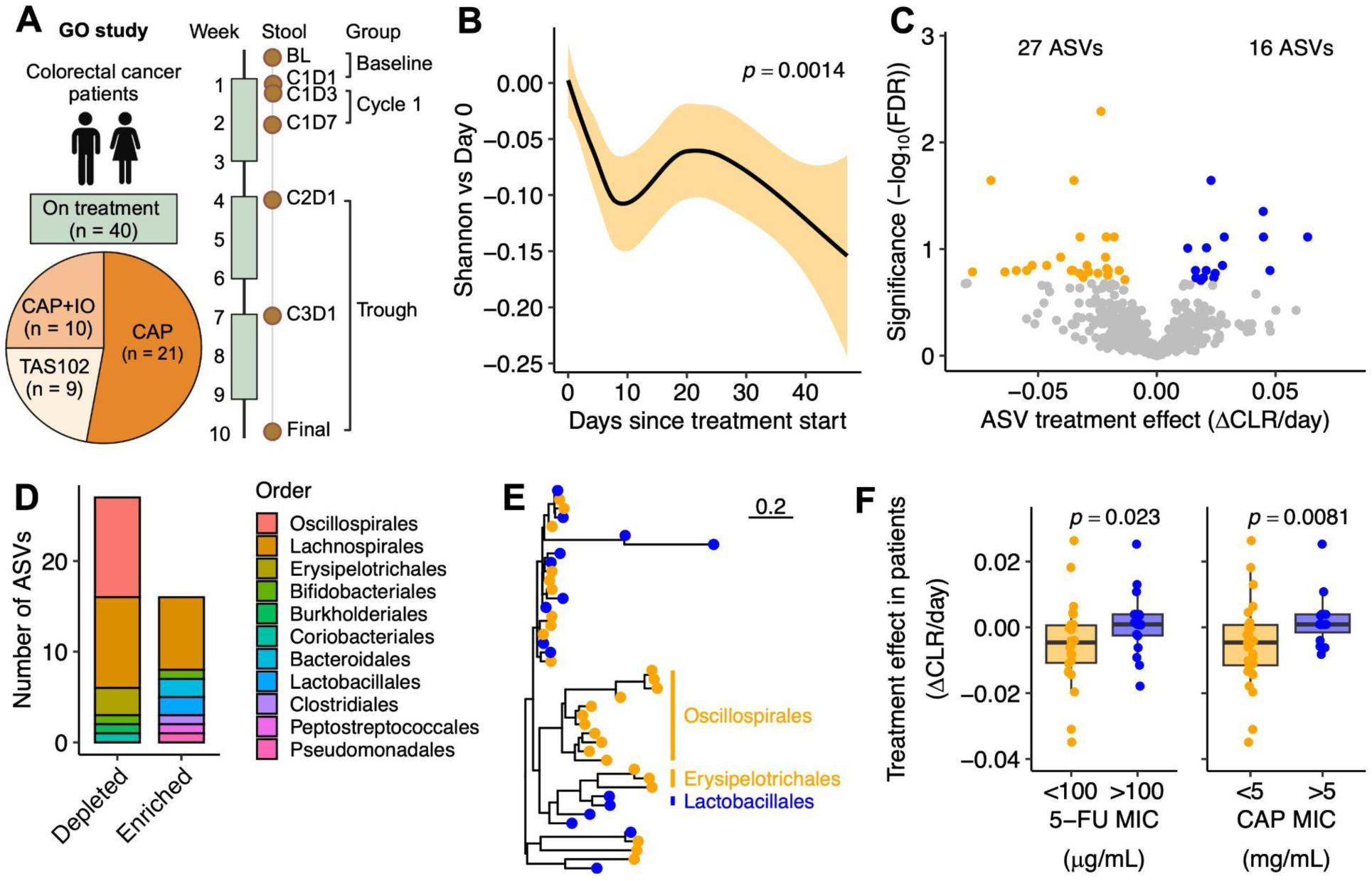
Oral fluoropyrimidine treatment impacts gut microbiome composition in colorectal cancer patients. **(A)** Gut microbiome and Oral fluoropyrimidine (GO) study design depicting patients treated with capecitabine (CAP), TAS-102, or combination CAP + immunotherapy (IO). Created with BioRender.com. **(B)** Change in alpha diversity during the study, normalized to baseline. *p*-value: mixed-effects model, Shannon ∼ Day + 1|Patient. Solid line with shading represents LOESS interpolation mean±sem. **(C)** Volcano plot of microbial amplicon sequence variants (ASVs) with respect to treatment time. *p*-value: mixed-effects model, central log ratio (CLR)-transformed Abundance ∼ Day + 1|Patient. Points represent enriched (blue) and depleted (orange) ASVs after treatment (false discovery rate (FDR) < 0.2). **(D)** Enriched and depleted ASV Order. **(E)** Phylogenetic tree of significantly differentially abundant ASVs, with labels for clades where treatment affected all clade members similarly (enriched [blue] or depleted [orange]). **(F)** Relationship between the impact of fluoropyrimidines on ASVs in patients and *in vitro* [minimum inhibitory concentrations (MICs) from (*16*)]. *p*-value: one-sided Mann-Whitney U test.

We utilized 16S rRNA gene sequencing to assess longitudinal shifts in the gut microbiota during cancer therapy. To better power our analysis, we sought to capture the overall longitudinal trends across all patients in all sub-cohorts. The combined dataset represented 217 samples from 40 patients with 67,915±3,744 high-quality reads per sample (**Data File S1**). Treatment was significantly associated with microbiota composition (**Fig. S2A**). For linear modeling, we truncated the data at 50 days post-treatment initiation to decrease the influence of long-duration outliers and patient drop-out on model results (**Fig. S1B**). Alpha diversity and amplicon sequence variants (ASVs) were analyzed using a mixed-effects model, with time as a fixed effect and Patient ID as a random effect (see *Methods*).

We observed a significant decrease in Shannon diversity index on treatment (**Fig. 1B**). Alternative markers of diversity trended lower but did not reach statistical significance (**Figs. S2B,C**). We identified 43 differentially abundant ASVs, including 16 that increased and 27 that decreased over time (**Fig. 1C, Fig. S2E, Data File S2**). The shifts in ASV abundance during Cycle 1 (samples from Cycle 1 Day 3 through Cycle 1 Day 14) and Trough (samples from Cycle 2 Day 1, Cycle 3 Day 1, or post-treatment) relative to baseline were correlated **(Fig. 1A, Fig. S2D)**. Treatment-associated ASVs were from 11 bacterial orders (**Fig. 1D**). The depleted ASVs included a cluster of *Oscillospirales,* whereas the enriched ASVs were more phylogenetically dispersed (**Fig. 1E**). Remarkably, our list of treatment-associated ASVs was significantly associated with previously reported values for bacterial sensitivity to CAP and 5-FU in mono-culture (*16*) (**Fig. 1F**), suggesting that at least some observed changes were due to the direct impact of fluoropyrimidines on the gut microbiota.

We investigated the impact of each individual treatment regimen on the microbiota. Baseline microbiota composition did not differ significantly by sub-cohort (PERMANOVA *p*>0.05, CLR-Euclidean ordination of ASV abundances). ASVs that were significantly altered in the global analysis were similarly altered across each sub-cohort (**Fig. S2F**). We detected significant enrichment of 15 ASVs and depletion of 19 ASVs in sub-cohort A alone (21 patients, **Data File S3**). We detected 4 significantly altered ASVs in sub-cohort B (9 patients, **Data File S3**) and no significantly altered ASVs in sub-cohort C (10 patients, **Data File S3**). Re-analysis of 9 patients randomly sampled from sub-cohort A revealed only four significantly altered ASVs (**Data File S3**), indicating that the lower number of differentially abundant taxa in sub-cohorts B and C are likely due to insufficient power.

We sought to study the impact of CAP on the microbiome in an independent patient cohort. We obtained previously published (*17*) 16S rRNA gene sequencing data from 33 CRC patients treated with CAP in the Netherlands (NE dataset, **Fig. S3A**). We observed a decline in ASV number, Inverse Simpson index, and Shannon index on treatment (**Fig. S3B-D**). Possibly due to the lower sample size and/or later sampling time relative to fluoropyrimidine treatment, we only detected a single significantly decreased ASV, mapped to the *Anaerovoracaceae* (**Fig. S3E,F**). Consistent with the GO dataset, the shift in ASV abundance in Cycle 3 was similar at Trough relative to Baseline (**Fig. S3G**). Despite many methodological explanations for a cohort effect, ASV-level shifts with respect to treatment were significantly associated between the GO and NE studies (**Fig. S3H**). Thus, it is possible to identify consistent shifts in the gut microbiota in CRC patients treated with oral fluoropyrimidines.

### Chemotherapy selects for gut microbial pyrimidine metabolism genes

Given the numerous potential mechanisms through which the gut microbiome can influence cancer progression and drug response, we sought to use metagenomic sequencing to develop a mechanistic hypothesis as to the consequences of fluoropyrimidine-induced shifts in the gut microbiome for drug response. Our metagenomic GO dataset represents 223 stool samples from 40 patients, with 25.9±0.7 million high quality reads per sample (7.67±0.22 Gbp, **Data File S4**). The distance to baseline based on metagenomic KEGG Orthologous group (KO) abundances (central log ratio (CLR)-Euclidean ordination) spiked during Cycle 1 with subsequent partial recovery (**Fig. S4A**); therefore, we opted to focus on KO abundances in Cycle 1 versus Baseline. Consistent shifts in KO abundance were detectable by principal coordinate analysis (**Fig. 2A**), reflecting 152 KOs with significantly altered relative abundance (**Fig. 2B, Data File S5**). Notably, nearly all (149) of these KOs increased in abundance, except for 3 KOs that decreased: K06989 (aspartate dehydrogenase), K20626 (lactoyl-CoA dehydratase subunit alpha), and K20461 (lantibiotic transport system permease protein). Gene set enrichment analysis of the 149 enriched KOs highlighted multiple pathways relevant to fluoropyrimidine metabolism, including pyrimidine metabolism, pantothenate and CoA biosynthesis, and β-alanine metabolism (**Fig. 2C**).

**Figure 2:**
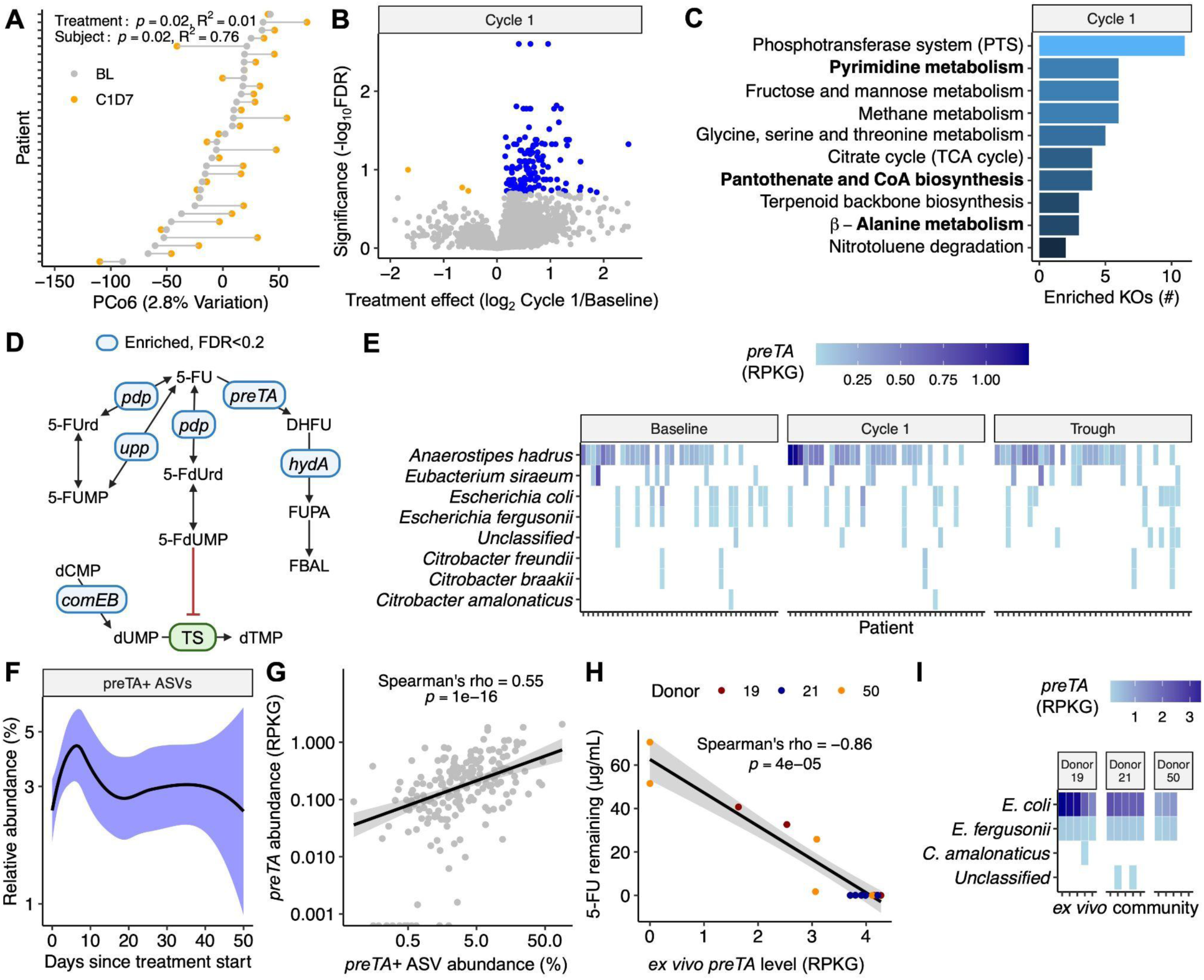
Oral fluoropyrimidine treatment selects for the bacterial drug metabolism gene *preTA*. **(A)** Reproducible shifts in gut microbial KEGG Ortholog (KO) abundance are observed across study participants with Baseline (BL) and C1D7 samples (*n* = 32) on the sixth principal coordinate (PCo6) of Euclidean distances between central log ratio (CLR)-transformed RPKGs. If a C1D1 but no BL sample was available, C1D1 was used instead. *p*-value: PERMANOVA with Patient ID as stratum. **(B)** Volcano plot of KO gene families with respect to treatment time for all *n* = 40 patients. Points represent significantly enriched (blue) and depleted (orange) gene families (FDR < 0.2). **(C)** Gene set enrichment analysis of significantly enriched KOs from **(B)**. Gene sets with *p* < 0.1 are displayed. Fluoropyrimidine metabolism-related pathways are bolded. **(D)** Fluoropyrimidine metabolism pathway with significantly enriched pyrimidine metabolism genes labeled in blue and 5-FU target thymidylate synthase (TS) labeled in green, created with BioRender.com. **(E)** Microbes contributing to *preTA* in patient samples during treatment, with patients ordered by mean *preTA* abundance. **(F)** Sum of preTA^+^ *Escherichia*, *Anaerostipes*, *Eubacterium, Citrobacter* amplicon sequence variant (ASV) relative abundance by 16S rRNA gene sequencing. Solid line with shading represents LOESS interpolation mean±sem. **(G)** Comparison of *preTA^+^* ASV abundance and *preTA* gene abundance. *p*-value: Spearman’s rank correlation. **(H)** *preTA* levels are associated with 5-FU depletion in *ex vivo* communities incubated with 50 µg/mL 5-FU for 48 hours anaerobically. *p*-value: Spearman’s rank correlation. **(I)** Microbes contributing to *preTA* in *ex vivo* communities. For **(G,H)**, solid lines represent a linear regression best fit with a shaded 95% confidence interval.

Inspection of the KOs within these pathways revealed increases in pyrimidine metabolism genes involved in fluoropyrimidine bioactivation, 5-FU clearance, and DNA biosynthesis (**Fig. 2D, Fig. S4B, Fig. S5**). Abundance of uracil phosphoribosyltransferase (*upp,* K00757) was significantly increased; *upp* is a well-established contributor to bacterial drug sensitivity (*18, 19*). Consistent with this trend, levels of pyrimidine-nucleoside phosphorylase (*pdp,* K00756) increased following the initiation of treatment. In turn, multiple genes in the bacterial pathway for 5-FU inactivation were enriched, including the bacterial dihydropyrimidine dehydrogenase (*preT,* K17722 and *preA*, K17723) responsible for the reduction of 5-FU to DHFU and dihydropyrimidinase (*hydA,* K01464) that processes DHFU to α-Fluoro-β-ureidopropionic acid. We further validated enrichment of pyrimidine metabolism genes using UniRef90 and enzyme class (EC) annotation schemes (**Figs. S4C-F**).

Given the clear clinical implications of the inactivation of 5-FU to DHFU (*11, 12*) and our prior work on the *preTA* operon (*16*), we further analyzed the bacterial taxa that explain the observed treatment-associated enrichment in *preTA* levels. Analysis of stratified gene abundance data revealed that *Anaerostipes hadrus*, *Escherichia* spp., *Eubacterium siraeum*, and *Citrobacter* spp. accounted for >99% of *preTA* **(Fig. 2E)**. Consistent with the observed inter-individual heterogeneity in the dominant *preTA^+^* species, no single ASV from these species was significantly enriched during Cycle 1 (false discovery rate > 0.2, mixed-effects model with Treatment Group as a fixed effect and Patient as a random effect). In contrast, the aggregated abundance of all *preTA*^+^ ASVs significantly increased following fluoropyrimidine treatment in both the GO (**Fig. 2F**) and NE (**Fig. S3I**) cohorts. Our aggregate metric for *preTA*^+^ ASV abundance by 16S rRNA gene sequencing was significantly associated with *preTA* gene abundance by metagenomics in the patient dataset (**Fig. 2G**).

To assess the impact of CAP on the gut microbiome along the intestinal tract, 500 mg/kg CAP was administered by oral gavage to 7-week-old mixed-sex C57BL/6J mice (*n* = 5-6/group) twice daily for two weeks (**Table S2**). 16S rRNA gene sequencing was performed on baseline stool and endpoint gastrointestinal contents (**Fig. S6A**). We observed significant differences in the mouse gut microbiota by body site (**Fig. S6B**), consistent with prior literature (*20*). Within each body site, there were significant differences between vehicle- and CAP-treated mice (**Fig. S6C**). Similar genus-level shifts were noted across body sites (**Fig. S6D**), including significant correlations between changes in stool relative to both ileum (**Fig. S6E**) and jejunum (**Fig. S6F**). Notably, we observed depletion of *Coriobacteriaceae* (**Fig. S6G**), enrichment of *Tyzzerella* (**Fig. S6H**), and enrichment of *preTA*-positive species across body sites (**Fig. S6I**). Thus, both the small and large intestinal gut microbiota are altered during treatment of mice with oral fluoropyrimidines.

We further validated these findings by testing the functional relevance of shifts in the abundance of *preTA* for 5-FU metabolism. Baseline stool samples from 3 patients with variable baseline *preTA* were cultured *ex vivo* in the presence of vehicle, 5-FU, or CAP for 48 hours. Both 5-FU and its oral prodrug CAP significantly inhibited overall bacterial growth (**Fig. S7A**), consistent with prior evidence for CAP bioactivation by human gut bacteria (*5*). 16S rRNA gene sequencing of the endpoint microbiotas revealed decreased diversity in response to CAP and 5-FU (**Fig. S7B**) and significantly altered community structure (**Fig. S7C**). The shifts in ASV abundance in response to CAP and 5-FU were significantly associated (**Fig. S7D**). We subjected post-incubation 5-FU-treated communities to metagenomic sequencing to measure *preTA* levels and LC-MS/MS to measure 5-FU. The quantity of 5-FU remaining varied between *ex vivo* communities and was significantly associated with *preTA* abundance (**Fig. 2H**). *E. coli* was the primary source of *preTA ex vivo* (**Fig. 2I**).

### Microbiota depletion exacerbates CAP toxicity

These data raised the intriguing hypothesis that an expansion of *preTA*^+^ bacteria following the initiation of fluoropyrimidine treatment could potentially mitigate gastrointestinal (GI) side effects. However, there was no established mouse model of oral fluoropyrimidine toxicity, necessitating extensive experimentation to establish such a model. We ultimately selected a daily mouse dose of 1,500 mg/kg CAP, which is metabolically equivalent to a daily human dose of 4,500 mg/m^2^ (*21*) and slightly higher than total daily GO study dose (1700-2500 mg/m^2^).

CAP was administered by oral gavage to 8-week-old female C57BL/6J mice (*n* = 8-12 mice/group; **Fig. 3A, Table S2**). CAP led to significant weight loss relative to vehicle controls (**Figs. 3B,C**) without any significant differences in body composition (**Figs. S8A-C**), small intestine length (**Fig. S8D**), or spleen weight (**Fig. S8E**). We detected clear evidence of colonic inflammation, including decreased colon length (**Fig. 3D**) and increased stool levels of lipocalin-2 (**Fig. 3E**). Hind paw thermal hyperalgesia was significantly worsened (**Fig. 3F**), indicating that this mouse model mimics aspects of the CAP-induced hand-foot syndrome (*22*), a common reason for CAP dose modifications (*8*). The severity of these phenotypes was significantly associated (**Fig. 3G**), motivating the use of weight loss as a proxy in subsequent experiments.

**Figure 3:**
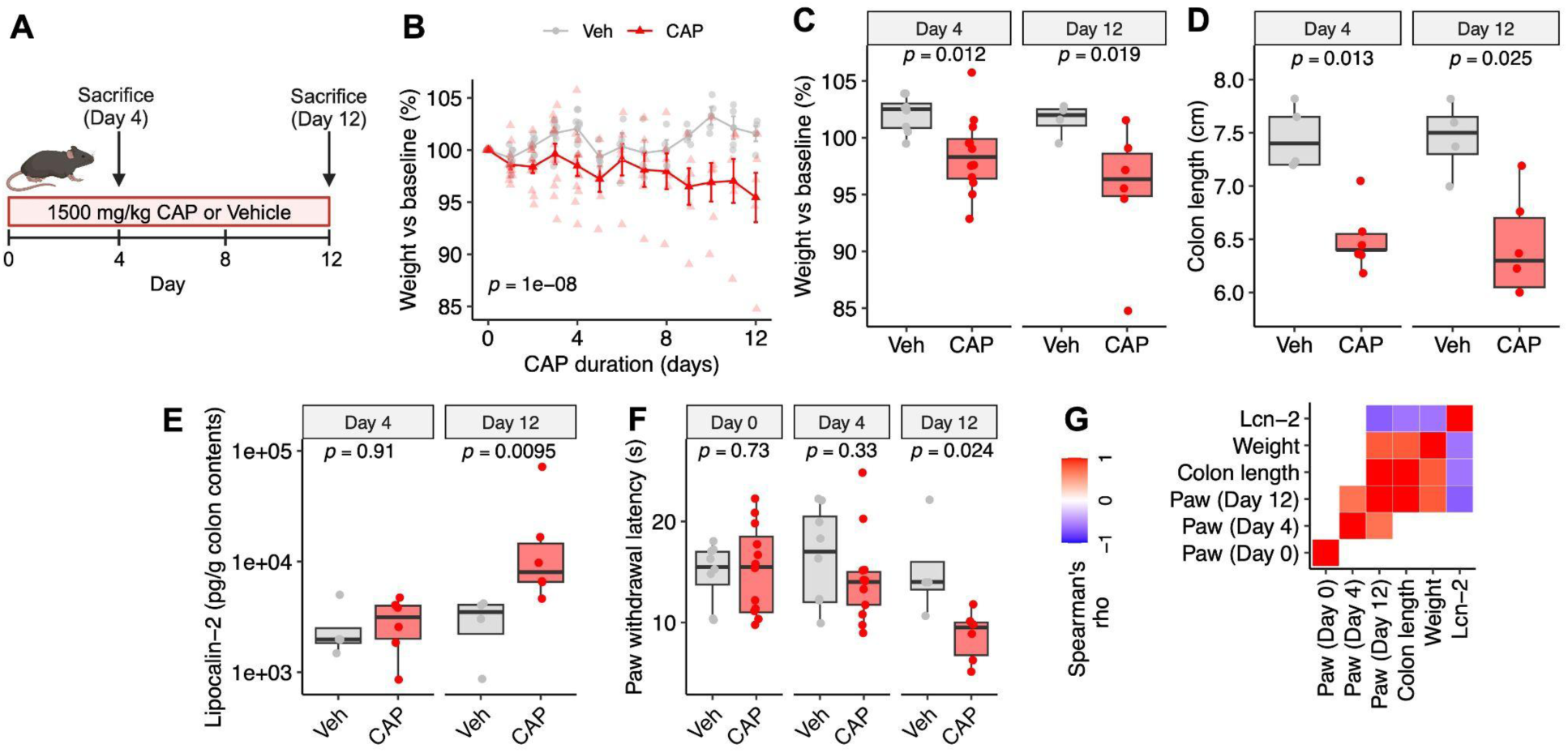
A mouse model of capecitabine (CAP) toxicity. **(A)** CAP toxicity mouse model (created with BioRender.com). Mice were gavaged daily with 1,500 mg/kg CAP (*n* = 12) or Vehicle (Veh, *n =* 8), with half the cohort sacrificed at Day 4. **(B)** Weight loss in CAP and Veh-treated mice. *p*-value: two-way ANOVA. **(C-F)** Toxicity endpoints at Day 4 and Day 12 of CAP treatment: mouse weight **(C)**, colon length **(D)**, lipocalin-2 in colon contents **(E)**, and hind paw withdrawal latency when placed on a 52°C hotplate **(F)**. *p-*values: Mann-Whitney U test. **(G)** Correlation of toxicity endpoints. All colored squares are significant (Spearman’s correlation *p*-value < 0.05).

CAP significantly altered the gut microbiota in our mouse toxicity model. We observed a significant decrease in Shannon diversity index (**Fig. S9A**). CAP treatment was significantly associated with microbiota composition (**Fig. S9B**). We identified 64 differentially abundant genera in CAP-treated mice and 0 in vehicle-treated mice, including 37 that increased and 27 that decreased over time (**Fig. S9C, Data File S6**). Treatment associated genera were from multiple bacterial orders, including many Clostridiales members **(Fig. S9D**).

Having established our mouse model, we tested if the overall gut microbiota impacts CAP toxicity. To deplete the gut microbiota, 6–8-week-old female C57BL/6J mice were treated with a cocktail of broad-spectrum antibiotics (ampicillin, vancomycin, neomycin, and metronidazole; AVNM) or vehicle for one week prior to CAP administration (**Fig. 4A, Table S2**). As expected, AVNM significantly decreased total colonization level and increased cecal weight (**Fig. S10A-B**). Weight loss was significantly accelerated in AVNM-treated mice (**Fig. 4B,C**), resulting in significantly decreased survival relative to vehicle controls (**Fig. 4D**).

**Figure 4:**
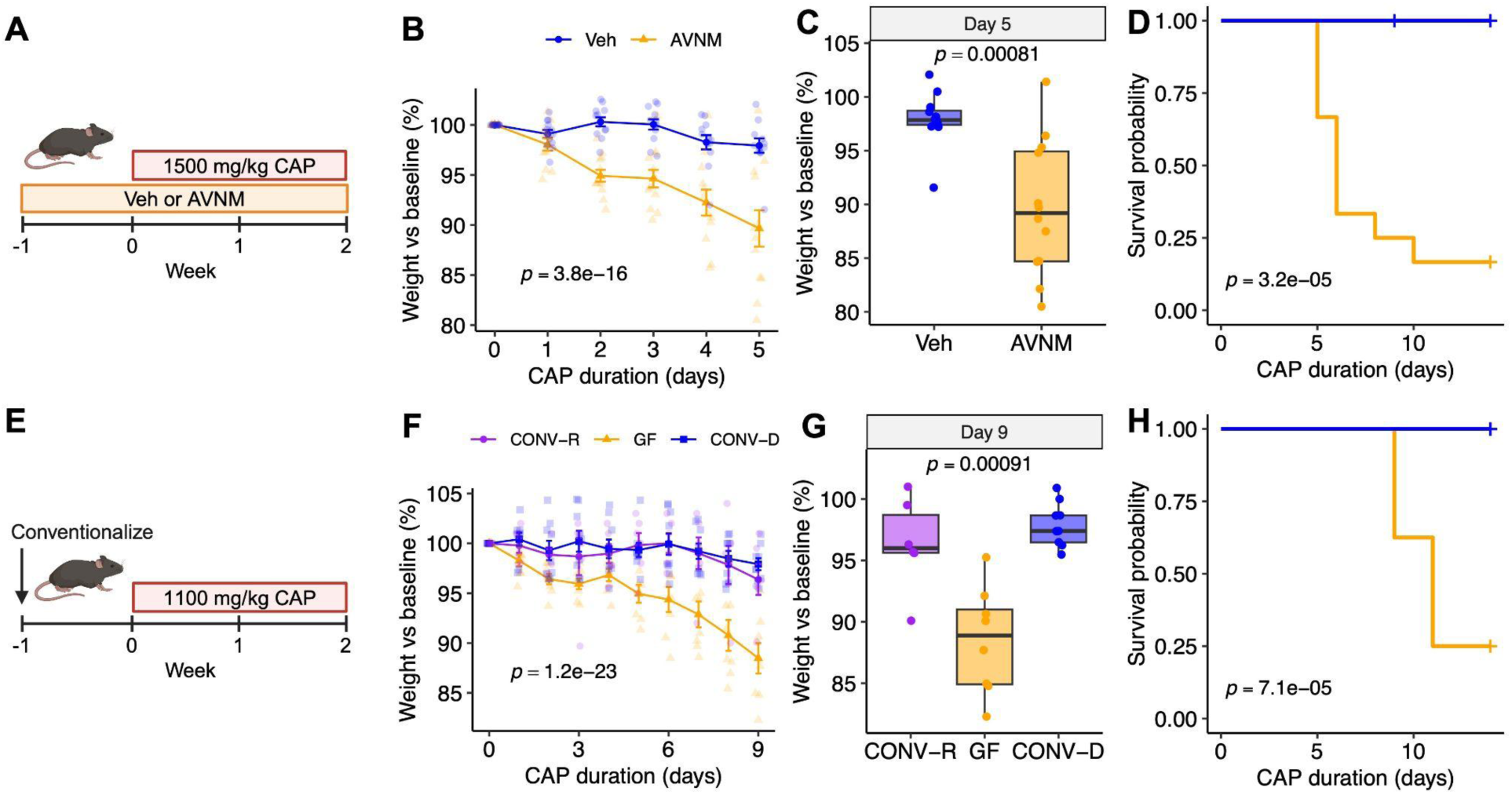
Microbiota depletion exacerbates capecitabine (CAP) toxicity in a mouse model. **(A)** AVNM antibiotic depletion CAP toxicity model (created with BioRender.com). Mice were treated with AVNM (*n* = 12) or Vehicle (Veh, *n* = 12) for 1 week, then gavaged daily with 1,500 mg/kg CAP. **(B)** CAP-induced weight loss in AVNM and Veh-treated mice. **(C)** Mouse weight on Day 5. **(D)** CAP survival curve in AVNM and Veh treated mice. **(E)** CAP toxicity model in conventionally reared (CONV-R, *n* = 8), germ free (GF, *n =* 8), and conventionalized (CONV-D, *n* = 8) mice. **(F)** CAP-induced weight loss in CONV-R, GF, and CONV-D mice. **(G)** Mouse weight on Day 9. **(H)** CAP survival curve in CONV-R, GF, and CONV-D mice. **(B,F)** *p-*values: two-way ANOVA **(B,F**), Mann-Whitney U test (**C**), Mantel-Cox test **(D,H)**, Kruskal-Wallis test (**G**).

These effects were validated in gnotobiotic mice. We compared CAP toxicity in germ-free (GF), conventionalized (ex-GF, CONV-D), and conventionally raised (CONV-R) 6–8-week-old female C57BL/6J mice (**Fig. 4E, Table S2**). Because GF mice experience higher serum 5-FU levels following CAP administration (*16*), we reduced the dose of CAP to 1,100 mg/kg by daily oral gavage. Consistent with our antibiotic depletion data, CAP led to significantly enhanced weight loss (**Fig. 4F,G**) and decreased survival (**Fig. 4H**) in GF mice relative to CONV-R or CONV-D mice. Taken together, these results indicate that an intact microbiota at the time of CAP treatment is critical to avoid increased drug toxicity.

### Microbial *preTA* rescues oral fluoropyrimidine toxicity

We leveraged this toxicity model to test the impact of bacterial *preTA* on CAP toxicity alone and in the context of a complex human gut microbiota. First, 6-8 week old mixed-sex C57BL/6J CONV-R mice were provided 5 g/L streptomycin in drinking water to enable the stable engraftment of isogenic Δ*preTA*, *preTA*-*wt* (*wt*), or *preTA* overexpressing (*preTA^++^*) *E. coli* MG1655 as described previously (*16*) (**Fig. 5A, Fig. S11A, Table S2**). *E. coli* colonization levels were similar between groups (**Fig. S10C, Fig. S11B**). Remarkably, the *preTA^++^* strain was sufficient to rescue CAP-induced survival deficits **(Fig. 5B, Fig. S11C)**, weight loss (**Fig. 5C, Fig. S11D-E**), and colon length (**Fig. 5D**) relative to the Δ*preTA* strain. The *preTA*-*wt* strain displayed an intermediate phenotype (**Fig. 5A-D**), consistent with its lower *preTA* expression in pure culture and strep-model colonized mice compared to *preTA*^++^ (**Fig. S12)**. The *preTA^++^*rescue effect was observed in both male and female mice (**Fig. S13**) and was validated in mono-colonized male mice (**Fig. S14**).

**Figure 5:**
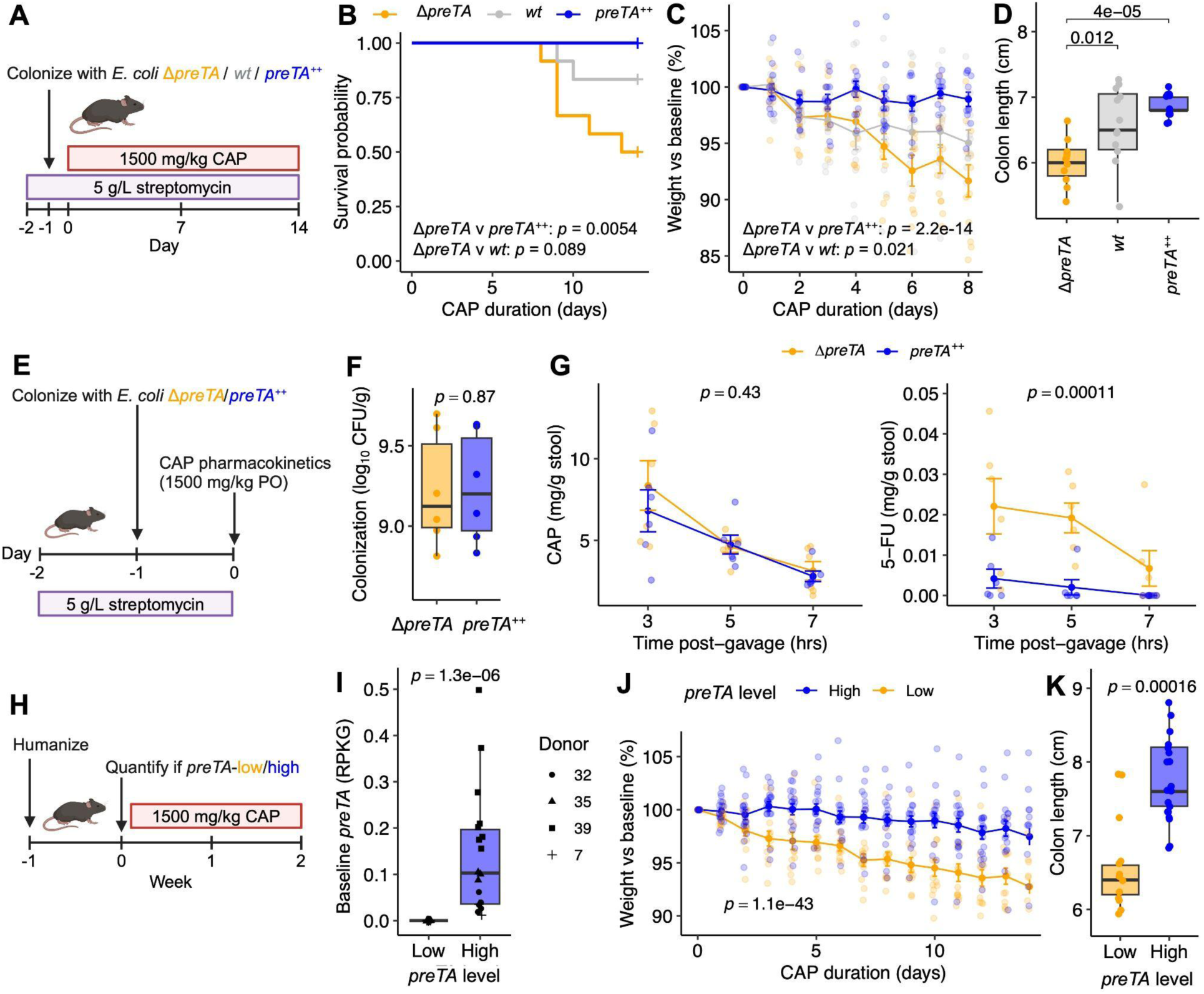
Bacterial *preTA* rescues CAP toxicity. **(A)** *preTA* streptomycin (strep) colonization CAP toxicity model. Mixed-sex mice were treated with strep for 1 day, gavaged with *E. coli* Δ*preTA* (*n* = 12), *wt* (*n* = 12), or *preTA*^++^ (*n* = 12), then gavaged daily with 1,500 mg/kg CAP. **(B)** CAP survival curve in *preTA* strep colonization model. **(C)** CAP-induced weight loss in *preTA* strep colonization model. **(D)** Colon length at sacrifice. **(E)** *preTA* strep colonization stool pharmacokinetics. 12 mice were treated with strep for one day, colonized with *E. coli* Δ*preTA* (*n* = 6) or *preTA*^++^ (*n* = 6) for one day, then given a single oral gavage of 1,500 mg/kg CAP with stool collection at 0, 3, 5, and 7 hours post-gavage. **(F)** *E. coli* colonization level at 0 hours post-gavage. **(G)** LC-MS/MS quantification of stool CAP and 5-FU levels at 3, 5, 7 hours post-gavage. **(H)** *preTA* humanization CAP toxicity model. 32 mice were colonized with stool from GO patients for 1 week (*n* = 8/donor), then gavaged daily with 1500 mg/kg CAP. **(I)** Baseline *preTA* level in humanized mice and group assignments. **(J)** CAP-induced weight loss in *preTA* humanization CAP toxicity model. **(K)** Endpoint colon length. *p*-values: Mantel-Cox test **(B)**, Mann-Whitney U test (**D,F,I,K**), group term from two-way ANOVA (**C,G,J**). **A,E,H** created with BioRender.com.

Next, we set out to measure direct metabolism of 5-FU (the active metabolite of CAP) in the intestinal tract. 9-week-old female C57BL/6J CONV-R mice were given 5 g/L streptomycin in drinking water, colonized with Δ*preTA* or *preTA^++^ E. coli*, and orally gavaged with 1,500 mg/kg CAP, followed by stool collection at 3-, 5-, and 7-hours post-gavage (**Fig. 5E, Table S2)**. *E. coli* colonization levels were similar between groups (**Fig. 5F**). CAP was detected at similar levels in both groups, with less CAP detected at later time points (**Fig. 5G**). In contrast, 5-FU was detected at significantly lower levels in mice colonized with *preTA^++^ E. coli* (**Fig. 5G**).

Finally, we probed the impact of *preTA* on CAP toxicity in a more complex community setting. We colonized GF 6–8-week-old male C57BL/6J mice with baseline stool samples collected from CRC patients in the GO study with varying *preTA* abundance (**Fig. 5H, Table S2**). We quantified *preTA* levels at baseline via metagenomic sequencing, finding measurable baseline *preTA* in 18/31 mice with evaluable stool (**Fig. 5I**). Consistent with our results with *E. coli*, mice with measurable baseline *preTA* experienced lower weight loss (**Fig. 5J**) and increased colon length (**Fig. 5K**).

We sought to assess the translational relevance of our findings in this mouse model of toxicity. We performed a retrospective chart review of oral fluoropyrimidine-associated side effects reported on the GO study. The majority (21/40) of GO subjects had at least one documented toxicity-related event, including 6 subjects with documented CAP dose reductions during the study (**Fig. 6A**). Consistent with our results in mice, *preTA* was significantly more abundant in patients without reported evidence of clinically significant toxicity (**Fig. 6B**). In addition to links to host drug toxicity, we noted a correlation between the magnitude of microbiota disruption and *preTA* abundance. Patients with high *preTA* maintained higher microbial diversity (**Fig. 6C**) and less deviation in microbial community structure relative to baseline (**Fig. 6D**). These trends were consistent across male and female patients (**Fig. S15**). We found no relationship between efficacy and *preTA* in our patient cohort; both baseline microbial KEGG Ortholog composition and baseline *preTA* levels were not significantly associated with disease progression (PERMANOVA *p*>0.05, **Fig. S16**, respectively).

**Figure 6:**
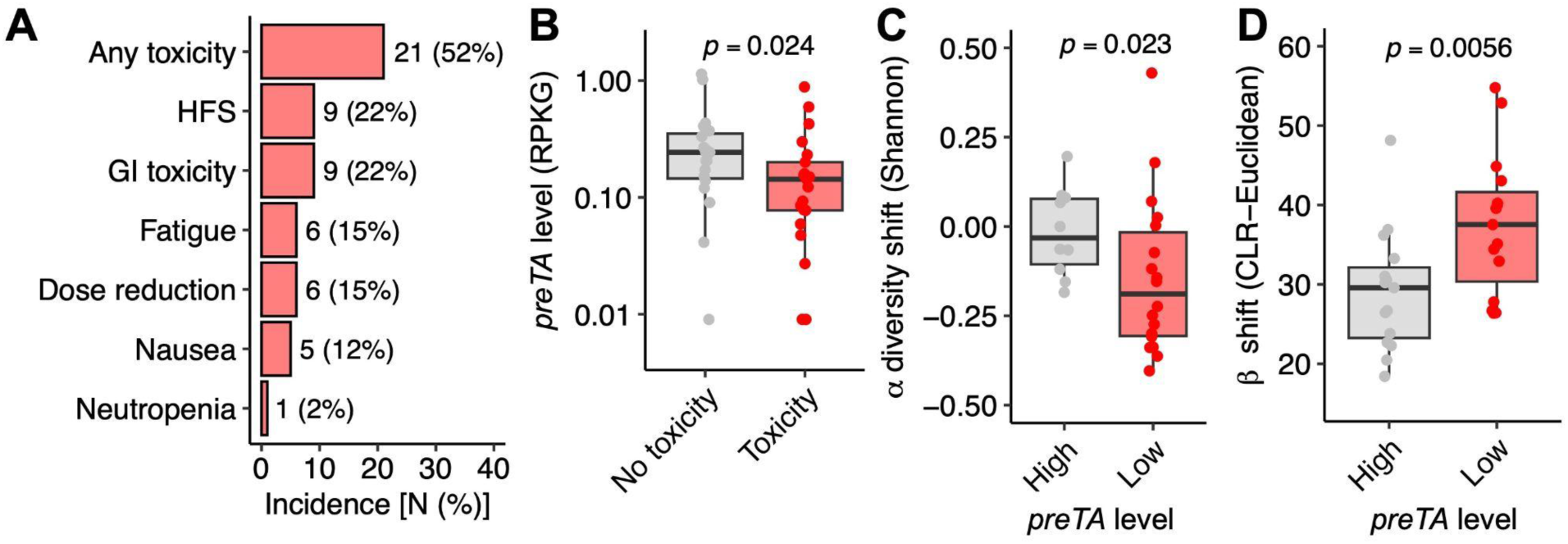
Bacterial *preTA* is associated with oral fluoropyrimidine toxicity in patients. **(A)** Distribution of toxicities in GO patients (*n* = 40). HFS: Hand-foot syndrome. **(B)** Patients experiencing toxicity have lower baseline stool microbial *preTA* levels (*n* = 40). *p*-value: one-sided Mann-Whitney U test. **(C)** *preTA* level (above/below mean) vs change in alpha diversity during Cycle 1 (C7D1-C1D1 Shannon index, *n* = 30). **(D)** *preTA* level (above/below mean) vs change in beta diversity during Cycle 1 (C7D1-C1D1 CLR-Euclidean distance, *n* = 30). **(C,D)** *p*-value: one-sided Wilcoxon signed-rank test.

We developed a model using baseline *preTA* to predict on-treatment toxicity using univariate received operating characteristic (ROC) curve analysis. Baseline *preTA* served as a mechanistic microbial predictor of toxicity with near-perfect class separation in the humanized mouse model (**Fig. S17A**). The area under the ROC curve (AUROC) was 0.68 when predicting any patient toxicity (**Fig. S17B**) and 0.74 for predicting dose reductions (**Fig. S17C**). To place these data in context of other genes, we calculated the AUROC for all KOs vs 5% weight loss (mice) or any toxicity (humans); *preTA* was in the top 1% of KOs for both mouse data (**Data File S7**) and human data (**Data File S8**). Notably, top-performing KOs included pyrimidine-related genes *preTA* homologue dihydroorotate dehydrogenase (K00254/K02823); dTMP kinase *tmk* (K00943); pyrimidine operon attenuation protein/uracil phosphoribosyltransferase *upp* (K00761/K02825); and 5’-nucleotidase *ushA* (K11751) (**Data Files S7,S8**).

## DISCUSSION

This study underscores the complex and clinically meaningful interplay between the gut microbiome and oral fluoropyrimidines, a commonly used chemotherapy. Longitudinal analyses of patient stool showed that chemotherapy has broad impacts on human gut microbial community structure, gene abundance, and functional pathways. This includes an enrichment for the bacterial *preTA* operon, which enables the inactivation of 5-FU to DHFU. We established and implemented a mouse model of CAP toxicity, allowing us to demonstrate that CAP toxicity is microbiome-dependent and that high levels of *preTA* expression are sufficient to control CAP toxicity in both male and female mice. Together, these findings emphasize the utility for the “reverse translation” of observations from cancer patients to gain mechanistic insight from follow-on mouse and cell culture experiments (*23, 24*).

Remarkably, we were able to detect consistent oral fluoropyrimidine-associated shifts in the gut microbiota in distinct patient cohorts from the United States and the Netherlands. This is particularly surprising given the geographical differences in the human gut microbiota (*25*) and prior literature emphasizing the difficulties in finding consistent differences across cohorts for cancer immunotherapy (*26*). While CAP altered the gut microbiome from jejunum to distal colon in our mouse model, future studies should profile human small intestinal contents during treatment by leveraging emerging spatial sampling technologies (*27, 28*). The longitudinal nature of frequent sampling is a strength of our approach, allowing us to assess changes in the microbiome over time by controlling for within-subject differences that confound cross-sectional studies. These results suggest that oral chemotherapy has a robust and reproducible impact on the human gut microbiota due to its ability to directly alter bacterial growth, as opposed to more indirect mechanisms at play for immunotherapy.

While we opted to focus on *preTA*, our results highlight numerous mechanisms through which chemotherapy-induced microbiome shifts could influence treatment outcomes. For example, we noted an overall decrease in microbial diversity following treatment, which has been associated with fluoropyrimidine-induced diarrhea in prior studies (*29*). Multiple bacterial taxa associated with immunotherapy response were depleted following chemotherapy, including the positively associated *Faecalibacterium prausnitzii* and negatively associated *Oscillospiraceae* species (*30, 31*). Notably, the most significantly enriched gene family was multidrug/hemolysin ABC transporter component *cylA* (K11050), a known virulence factor (*32*) that has also been implicated in inflammatory bowel disease (*33*). Thus, shifts in the microbiome following oral fluoropyrimidine treatment likely results in a complex combination of beneficial, neutral, and deleterious shifts in gut microbial ecology and host-microbiome interactions.

Our results emphasize the complex role the gut microbiome can play in chemotherapy response. In this study, we utilized a novel mouse model to demonstrate that CAP toxicity is microbiome-dependent, consistent with data in human subjects showing heightened toxicity of CAP-containing regimens in antibiotic-pretreated patients (*34*). On the other hand, extensive prior literature has shown that antibiotics alleviate the GI toxicity of the anti-cancer drug irinotecan (SN-38) in rats and humans by preventing bacterial reactivation of inactive SN-38G (*35, 36*). Since treatment regimens such as XELIRI and FOLFIRI combine both fluoropyrimidines and irinotecan (*37*), use of antibiotics may worsen CAP toxicity while alleviating irinotecan toxicity, highlighting a clinical need to develop targeted microbial manipulations rather than relying on broad-spectrum antibiotics.

The observation that bacterial *preTA* is sufficient to control CAP toxicity in mice has clear translational relevance, raising the potential to utilize current or next-generation probiotics to ameliorate GI toxicity in cancer patients. GI toxicities usually present early during CAP treatment (*38, 39*). This observation is consistent with decreased toxicity following *preTA* expansion during treatment. Notably, prior studies have shown a benefit of fecal microbiota transplantation (FMT) and probiotics containing *Lactobacillus reuteri* (*40, 41*), both of which could be in part driven by increased *preTA* and thus enhanced 5-FU inactivation within the gut lumen. Although CAP toxicities differ by sex in clinical practice (*42*), our findings demonstrate that *preTA* can modulate toxicity in both male and female mice. A critical next step is to assess the trade-offs between decreased toxicity and efficacy, especially given prior data suggesting that high levels of *preTA* can interfere with the anti-tumor effects of CAP (*16*). While *preTA* was associated with toxicity but not efficacy in the GO dataset, further work is needed to optimize probiotics to mitigate toxicity without diminishing efficacy prior to clinical translation. More broadly, probiotic strains that express *preTA* could be a valuable tool to treat pyrimidine-related toxicity in other conditions, potentially depriving cancer cells of uracil in pancreatic ductal adenocarcinoma (*43*) and clearing excess uracil in patients with clinical DPYD deficiency (*44*).

Low *preTA* levels were predictive of increased adverse effects in humanized mice and patients, providing a *proof-of-concept* for the use of microbial “genotyping” as a tool for predicting toxicity. Baseline *preTA* serves as a mechanistic microbial predictor of toxicity with near-perfect class separation in humanized *Dpyd*^+/+^ mice and an AUROC of 0.68 in the GO patient cohort. Meanwhile, clinical predictors in *DPYD*^+/+^ patients combining age, body surface area, type of treatment regimen, and creatinine levels achieve an AUROC of 0.68 when predicting global Grade 3+ toxicity (*45*), and comprehensive DPYD genotyping alone has an AUROC near 0.5 for predicting global Grade 2+ toxicity (*46*). While our AUROC < 1 suggests that additional mechanisms are at play and necessitates further integration with clinical variables and/or host genetics, our microbiome-based prediction algorithm could help address long-standing challenges in avoiding the severe toxicities associated with fluoropyrimidine drugs.

Our results highlight the role of the gut microbiome in the pharmacodynamics of orally bioavailable drugs. CAP is thought to be nearly 100% orally bioavailable; in an initial study of 6 cancer patients given a single ^14^C-labeled CAP dose, 95±8% of labeled metabolites were recovered from urine, with only 0.69-7% recovered in stool (*47*). In larger subsequent studies, urine metabolite recovery was lower and more variable (70±17%) (*48, 49*), consistent with higher fecal excretion. Nucleotide transporters including hCNT1, hCNT3, OATP, hENT1, and hENT2 allow CAP and its metabolites to move between circulation, gut tissue, and the gut lumen (*50–55*), enabling extensive gut microbiota-CAP interactions prior to renal clearance. Consistent with these prior studies, we detected both CAP and 5-FU in the stool of mice, with significantly lower stool 5-FU levels in mice colonized with *preTA* expressing *E. coli*. This local microbial control of intestinal drug levels could play a role in mitigation of gastrointestinal chemotoxicities. Future studies should seek to provide a more detailed characterization of the interaction between host and microbial metabolism along the gastrointestinal tract and in other organs like the liver.

While this work reinforces the crucial role of drug-microbiome interactions in cancer therapy, key translational questions remain. Heterogeneous patient populations, combinatorial treatment regimens, and small sub-cohort sample sizes limited our ability to test the impact of immunotherapy, radiation, and biologics on the microbiome. Though we observed similar taxonomic shifts across all three cohorts (CAP, TAS102, and CAP plus pembrolizumab), better-powered studies are needed to understand how concurrent treatments modulate CAP-microbiome interactions. While severe diarrhea is a common side effect in patients (*56*), we did not observe diarrhea in our mouse model, potentially due to variation in metabolic rates and/or physiological toxicities between mice and humans (*21, 57*). Since host DPYD deficiency is associated with life-threatening side effects (*14*), future work seeking to understand toxicity should integrate genetics of both host *DPYD* and microbial *preTA*. Data in human subjects suggests that orally delivered fluoropyrimidines have ten times less neutropenia but comparable GI toxicity to infusional 5-FU (*58, 59*). The degree to which bacterial *preTA* is better able to mitigate toxicities following oral versus intravenous delivery is an important area for future studies.

Overall, our results highlight the complex and bidirectional relationship between the gut microbiome and oral fluoropyrimidine chemotherapy and the consequences of this relationship for host drug toxicity. Integrating longitudinal microbiome analysis with *in vivo* and *ex vivo* experiments, we show that fluoropyrimidine metabolism gene *preTA* is a critical human gut bacterial mediator of oral fluoropyrimidine toxicity. This microbial operon serves as a biomarker for predicting and a novel target for controlling dose-limiting toxicities. Our findings underscore the necessity of considering the microbiome as an integral component in the pharmacological landscape, paving the way for microbiota-based precision therapy.

## MATERIALS AND METHODS

### Study design

The <u>G</u>ut microbiome and <u>O</u>ral fluoropyrimidines (GO) study was an observational clinical study registered at ClinicalTrials.gov under the identifier NCT04054908. All participants provided informed consent. The study was approved by the UCSF Institutional Review Board. Participants were recruited and sampled, without any compensation, at UCSF from the study start date (2018/04/13) to end date (2022/06/30). Inclusion criteria included: (1) 18 years or older, (2) histologically confirmed colorectal adenocarcinoma, and (3) expected to receive oral fluoropyrimidine therapy. Patients who met any of the following criteria were excluded: (1) known HIV positive diagnosis, (2) chemotherapy, biologic or immunotherapy in the previous 2 weeks, (3) exposure to ≥2 weeks antibiotics in the last 6 months, (4) exposure to antibiotics in the past 4 weeks. Patients were enrolled to one of three cohorts: Cohort A received oral CAP as part of standard-of-care therapy; Cohort B received TAS-102 (trifluridine/tipiracil) ± Y-90 radioembolization (*60*); Cohort C received CAP + immunotherapy (pembrolizumab) + bevacizumab as part of a separate clinical trial (NCT03396926). Patients with concurrent rectal radiation therapy (RT) were originally excluded, but the eligibility criteria was amended to include patients with concurrent rectal RT to increase enrollment. CAP and TAS-102 tablets were prescribed according to FDA labels for oral dosing. CAP tablets were taken twice daily on days 1–14 of a 21-day Cycle (no RT) or on days of radiation only (RT); TAS-102 tablets were taken twice daily on days 1–5 and 8–12 of a 28-day Cycle. Stool collection occurred at home before chemotherapy initiation and on day 1 of Cycles 1, 2, 3. During Cycle 1, stool was also collected at day 3 and midpoint (day 7 for Cohorts A, C; day 10 for cohort B). Actual collection days are plotted in **Fig. S1**. Stool samples were collected on fecal occult blood test (FOBT) cards for all time points. Additional bulk scoop samples were collected at baseline for culturing. Stool was stored at −80 °C upon receipt at UCSF. 222 stool samples from 40 participants on FOBT cards were ultimately evaluable; 4 samples were not evaluable due to insufficient extracted DNA quantity. Methods for longitudinal sampling of small intestinal contents were not widely available at the time of GO study initiation (April 13th, 2018).

### Resources Table

| Reagent | Source | Catalog Number |
| --- | --- | --- |
| Capecitabine | Fisher Scientific | 50148375 |
| Capecitabine-$^2\text{H}_{11}$ | Santa Cruz Biotechnology | 1132662 |
| 5-fluorouracil | Millipore Sigma | F6627 |
| 5-fluorouracil-$^{13}\text{C}$, $^{15}\text{N}_2$ | Santa Cruz Biotechnology | 217189 |
| Citric acid | Thomas Scientific | C752W30 |
| Sodium citrate dihydrate | Fisher Scientific | 18613454 |
| Gum arabic | Spectrum Chemicals | GU113 |
| *Escherichia coli* $\Delta\text{preTA}$ | (16) | N/A |
| *Escherichia coli preTA-wt* | (16) | N/A |
| *Escherichia coli preTA<sup>++</sup>* | (16) | N/A |
| Bacto Brain Heart Infusion (broth) | Fisher Scientific | 237500 |
| L-Cysteine HCl | Fisher Scientific | C81020 |
| Hemin | Fisher Scientific | 51280 |
| Menadione (Vitamin K) | Fisher Scientific | 102259 |
| Lysogeny broth (LB) | Millipore Sigma | L3152 |
| Eosin Methylene Blue (EMB) agar | Thomas Scientific | C5701 |
| Streptomycin | Thomas Scientific | S6501 |
| Neomycin | Neta Scientific | SIAL-N1876 |
| Azithromycin | Neta Scientific | FISHUCOP-A20765G |
| Vancomycin | Thomas Scientific | C752X68 |
| Metronidazole | Neta Scientific | SIAL-M1547-25G |
| Sucrose | Thomas Scientific | C752W82 |
| Dimethyl sulfoxide | Millipore Sigma | D8418 |
| Mouse Lipocalin-2/NGAL DuoSet ELISA | Fisher Scientific | DY185705 |
| ZymoBIOMICS 96 MagBead DNA Kit | Zymo Research | D4302 |
| SYBR Green I | Millipore Sigma | S9430 |
| Nuclease-free water | Life Tech | 0977-023 |
| Phusion High-Fidelity PCR Kit | Thermo Scientific | F553L |
| QIAquick Gel Extraction Kit | Qiagen | 28604 |
| KAPA Library Quantification Kit | Roche | KK4824 |
| MiSeq Reagent Kit v3 (600 cycles) | Illumina | MS-102-3003 |
| Illumina DNA Prep, (M)<br>Tagmentation (96 Samples, IPB) | Illumina | 20060059 |
| Quant-iT Picogreen dsDNA Kit | Invitrogen | P7589 |
| SYBR Select | Life Technologies | 4472908 |
| Wildtype C57BL/6J mice | Jackson Labs | 000664 |
| Standard chow diet (SPF) | LabDiet | 5058 |
| Standard autoclaved chow diet | LabDiet | 5021 |
| BreatheEasy Covers | Millipore Sigma | Z380059 |

### DNA extraction

ZymoBIOMICs 96 MagBead DNA Kit was used for DNA extractions. 750 µL of lysis solution was added to fecal aliquots (20-50 mg) in lysis tubes, while cell pellets from *ex vivo* communities were resuspended in 200 µL lysis solution and transferred to lysis tubes containing 550 µL lysis solution for a final volume of 750 µL lysis solution. Samples were homogenized with 5 min bead beating (Mini-Beadbeater-96, BioSpec), followed by 5 min room temperature (RT) incubation, and repeat 5 min bead beating. Samples were centrifuged for 1 min at 15,000 rcf, with 200 µL supernatant transferred into 1 mL deep-well plates and purified according to the manufacturer’s instructions.

### 16S rRNA gene sequencing

The V4 region of the 16S rRNA gene was amplified using primers targeting 515F/806R regions (**Table S3**). The reaction mix was 0.45 μL DMSO, 0.0045 μL SYBR Green I 10x diluted in DMSO to 1000x, KAPA HiFi PCR kit (1.8 μL 5x KAPA HiFi Buffer, 0.27 μL 10 mM dNTPs, 0.18 μL KAPA HiFi polymerase), 0.045 μL of each amplification primer (final concentration 1 μM), 6.2055 μL nuclease-free water, and 1 μL DNA. A BioRad CFX 384 real-time PCR instrument amplified four 10-fold serial dilutions of DNA with the following parameters: 5 min 95°C, 20x(20 sec 98°C, 15 sec 55°C, 60 sec 72°C), hold 4°C. Non-plateaued individual sample dilutions were selected for indexing PCR. KAPA HiFi PCR kit was used with: 4 μL 5x KAPA HiFi Buffer, 0.6 μL 10 mM dNTPs, 1 μL DMSO, 0.4 μL KAPA HiFi polymerase, 4 μL indexing primer, 10 μL of 100-fold diluted primary PCR reaction. Secondary PCR amplification parameters were identical to Primary PCR. Amplicons were quantified with PicoGreen according to manufacturer’s instructions, equimolar pooled, and gel purified (QIAquick Gel Extraction Kit). Libraries were quantified with KAPA Library Quantification Kit for Illumina Platforms according to the manufacturer’s instructions, spiked with 15% PhiX, and sequenced on an Illumina MiSeqV3 instrument. Primers and adapters were removed using the cutadapt trim-paired command in QIIME2 (v2020.11) (*61*). Sequences underwent trimming to 220 bp (forward) or 150 bp (reverse), quality filtering, denoising, and chimera filtering using dada2 (v1.18.0) with QIIME2 command denoise-paired (*62*). Sequence length was filtered to 250-255 bp with QIIME2 command feature-table filter-seqs. Taxonomy was assigned to amplicon sequence variants (ASVs) using the SILVA v138 database (*63*). Sequence variants not present in ≥3 samples with ≥10 reads were removed from downstream analysis.

### 16S rRNA gene analysis

Alpha diversity metrics were computed with Phyloseq command estimate_richness (*64*). Differential alpha diversity was calculated with mixed effects model Diversity ∼ Day + 1|Patient with the nlme (v3.1-164) command lme. ASV relative abundances were central log ratio (CLR)-transformed prior to further analysis. Principal coordinate analysis (PCoA) was performed with command prcomp (CLR-Euclidean ordination). PERMANOVA was performed using vegan (v2.6-4) commands vegdist (CLR-Euclidean ordination) and adonis2 (*65*), with Patient as a strata. Differential abundance was calculated using mixed effects model Abundance ∼ Day + 1|Patient, followed by false discovery rate (FDR) correction with R command p.adjust, with FDR < 0.2 called as significant. ASV phylogenetic tree construction was performed with QIIME2 command phylogeny align-to-tree-mafft-fasttree. Line plots were plotted using ggplot2 (v3.5.1) command geom_smooth with LOESS regression (*66*). For comparisons of patient sensitivity with *in vitro* sensitivity, strain minimum inhibitory concentrations (MICs) were taken from (*16*).

### Metagenomic sequencing

Shotgun libraries were prepared using the Illumina DNA Prep Tagmentation kit according to the manufacturer protocol. Libraries were assessed with PicoGreen and TapeStation 4200 (Agilent) for quantity and quality checks. Paired end libraries were sequenced using S1 flow cells on Illumina NovaSeq 6000 platforms. Demultiplexed reads underwent adapter trimming and quality filtering with FastP (v0.23.2) (*67*) and host read removal by mapping to human genome (GRCh38) with BMTagger (v3.101) (*68*). Genome equivalents were quantified with microbeCensus (v1.1.1) (*69*). Taxonomy was annotated with MetaPhlAn 4 (*70*). Genes were annotated with HUMAnN 3.0 (*71*), with UniRef90 families further mapped to KEGG Ortholog and Enzyme Class.

### Metagenomic analysis

Gene abundances were normalized to reads per kilobase per genome equivalent using microbeCensus values prior to downstream analysis (*69*). PCA was performed with command prcomp (CLR-Euclidean ordination). PERMANOVA was performed using vegan commands vegdist (CLR-Euclidean ordination) and adonis2 (*65*), with Patient as a strata. Differential abundance was calculated with mixed effects model Abundance ∼ Day + 1|Patient using the nlme command lme, followed by FDR correction, with FDR < 0.2 called as significant. Gene set enrichment analysis was performed on significant genes using clusterProfiler (v4.6.2) command enrichKEGG with a q-value cutoff of 0.2 (*72*). Line plots were plotted using ggplot2 command geom_smooth with LOESS regression (*66*).

### Generation of *ex vivo* communities

Three patients from Cohort A with variable *preTA* levels and Baseline vs C1D7 diversity were selected for *ex vivo* communities. Brain Heart Infusion supplemented with L-cysteine hydrochloride (0.05% w/v), hemin (5 μg/ml), and vitamin K (1 μg/ml), termed BHI+, was used as media. 100 mg baseline stool from each patient was thawed and added anaerobically to 1 mL BHI+, vortexed for 1 min, and incubated 5 min at RT to allow debris to settle. 500 µL supernatant (“fecal slurry”) was transferred to a new tube. 5-FU was dissolved in dimethylsulfoxide, supplemented at 1% (v/v) in BHI+ media and assayed at 50 ug/mL. 1% DMSO was added to BHI+ for CAP and vehicle groups; CAP was directly resuspended in this media and assayed at 5 mg/mL. 5 µL fecal slurry was inoculated in sextuplicate into 195 µL of media ± drug in a 96-well plate, with negative control wells to confirm media sterility. Plates were covered with BreathEasy covers and incubated at 37°C for 48 hr in a Gen5 plate reader, with 1 min linear shake prior to OD_600_ readings every 15 min. Plates were removed from plate reader and spun at 3000 rpm for 30 min, followed by supernatant (“spent media”) transfer to a new plate. The plate of cell pellets was frozen at −80°C for future DNA extraction followed by 16S rRNA gene (all samples) and metagenomic sequencing (5-FU-treated communities only).

### Extraction of fluoropyrimidine metabolites from *ex vivo* communities

Spent media was thawed on ice. 50 µL spent media was dissolved in 150 µL water and 800 µL organic phase (50% ACN, 50% MeOH). A standard curve was generated by dissolving serial dilutions of 5-FU in DMSO into BHI+. 10 µL of 50 ng/ml 5-fluorouracil-^13^C,^15^N_2_ internal standard was spiked into all samples. Samples were vortexed 5 min and incubated on ice for 30 min. Extraction mixture was spun down at 15000 rcf for 20 min. 250 µL extraction supernatant was dried in speed vac and resuspended in 200 µL of 10% ACN in water for injection.

### Extraction of fluoropyrimidine metabolites from stool

Stool was collected from mice and flash-frozen in liquid nitrogen, followed by lyophilization for 27 hours. Dry stool weight was recorded. Dry stool was then homogenized at 4C with the following settings: 3 x [20 sec 6400 rpm, 30 sec pause], followed by dissolution in 800 µL of 2:2:1 ACN:MeOH:water. A standard curve was generated by spiking in serial dilutions of CAP and 5-FU in MeOH into stool from untreated mice. 10 µL of 500 ng/ml capecitabine-^2^H_11_ and 50 ng/mL 5-fluorouracil-^13^C,^15^N_2_ in MeOH was spiked into all samples. Samples were vortexed 5 min and incubated on ice for 30 min. Extraction mixture was spun down at 15000 rcf for 10 min. 250 µL extraction supernatant was dried in the speed vac overnight and resuspended in 1000 µL of 10% ACN in water. This mixture was centrifuged at 15000 rcf for 10 minutes, with 100 µL supernatant taken for injection.

### LC-MS/MS quantification of CAP and 5-FU from stool

Quantification was performed using a validated fluoropyrimidine quantification protocol (*16*). Samples were loaded into a Synergi column on a SCIEX Triple Quad 7500 instrument with a linear ion QTRAP. Chromatographic separation was achieved using a Phenomenex Synergi column 4 µM Fusion RP-80 (50 × 2 mm) at 35 °C. The mobile phase was methanol + 0.1% formic acid (A) and HPLC-grade water + 0.1% formic acid (B). A flow rate of 0.4 ml/min was used for the following gradient elution profile: 0% B at 0-2 min, gradient to 100% B from 2 to 5.9 min, gradient to 0% B at 5.9-6 min. The autosampler was maintained at 4 °C. Eluate from the column was ionized in the LC-MS/MS using an electrospray ionization source in positive polarity. Peak areas were calculated using built-in SCIEX OS software. For quantification from *ex vivo* communities, 5-FU peak area was normalized to the internal standard, with concentration calculated based on the 5-FU standard curve (Pearson R^2^ = 1.00). For quantification from stool, 5-FU and CAP peak areas were normalized to the internal standard, with concentration calculated based on the stool spike-in 5-FU and CAP standard curves (Pearson R^2^ = 1.00 and 0.97, respectively).

### Reverse-transcription quantitative polymerase chain reaction (RT-qPCR) quantification of *preTA* from culture and mouse stool

*E. coli* MG1655 strains (*E. coli* Δ*preTA*, *wt*, *preTA*^++^) used were generated previously (*16*). For RNA extraction from culture, strains were cultured in quintuplicate in BHI overnight, and pelleted at 15,000g. For mouse colonization and RNA extraction from mouse stool, streptomycin water was prepared from autoclaved tap water and supplemented with 5 g/L streptomycin, followed by 0.2 μm filter sterilization. 12 mice were treated *ad libitum* with streptomycin water for the duration of the experiment. Overnight cultures of engineered *E. coli* strains (*E. coli* Δ*preTA*, *wt*, *preTA*^++^) were pelleted by centrifugation, washed with an equal volume of sterile 0.85% saline, pelleted by centrifugation, and resuspended in 1:10 sterile saline. Mice were gavaged with 200 µl bacterial suspension (4 Δ*preTA*, 4 *preTA-wt*, 4 *preTA*^++^) by gavage following 1 day on strep water. One day post-colonization, stool was collected. Pellets and stool were (re)suspended in 500 μL TRI reagent in Lysing Matrix E tubes, subjected to two rounds of 5 min bead beating, and stored at −80°C. RNA was purified with the Direct-Zol RNA miniprep kit according to the manufacturer protocol and quality-controlled with NanoDrop (A260/A280 = 1.87-2.09, A260/A230=1.92-2.57). cDNA was transcribed using the QuantiTect Reverse Transcription kit per manufacturer protocol. Primers for *preTA* and housekeeping gene *rrsA* were validated and optimized with gradient PCR (**Table S3**). cDNA was serially diluted (1-1000x) in RNAse-free water to remove PCR inhibitors. Each well contained 2 μL cDNA template and 8 uL SYBR master mix with primers (ratio of 1 μL 10 uM forward and reverse primers, 5 μL 2x SYBR Green Master Mix, 2 uL RNAse-free water) per well of the 384-well plate. The following qPCR reaction was run: 2 min 50°C, 2min 95°C, 40x(15 sec 95°C, 1 min 60°C), melt curve from 65 to 95°C. Relative expression was calculated using the delta-delta Ct method (*73*).

### Mouse studies

All animal experiments were conducted under protocol AN200526 approved by the UCSF Institutional Animal Care and Use Committee. GF C57BL/6J mice were born at UCSF; specific pathogen-free (SPF) C57BL6/J mice were purchased from Jackson Laboratory. Diets were provided *ad libitum*, with standard chow diet (LabDiet 5058) in the SPF facility and standard autoclaved chow diet (LabDiet 5021) in the gnotobiotic facility. Mouse age, sex, and caging is reported in **Table S2**. Mice were housed at temperatures ranging from 67 to 74°F and humidity ranging from 30 to 70% in light/dark cycle 12h/12h. GF mice were maintained within the UCSF Gnotobiotic Core Facility. Stool pellets from the Gnotobiotic Core Facility were screened every 2 weeks by culture and qPCR, whenever mice were transferred between isolators, and at the beginning and end of each experiment using V4_515F_Gnoto/V4_806R_Gnoto universal primers (**Table S3**). In CONV-R mice, cage swaps were performed 1 week before experiments to minimize confounding cage-dependent microbiota effects (equal numbers of mice from each original cage were separated into experimental cages split across experimental groups). To enable these cage swaps, we used predominantly female mice (**Table S2**), which are less likely to fight following changes in their cage mate (*74*). In the gnotobiotic facility, we used predominantly male mice based on germ-free mouse availability (**Table S2**).

### CAP dose selection

We performed a human-to-mouse dose conversion to account for metabolic differences (from (*21*)):

2500 mg/m^2^ human dose x 1/37 m^2^/kg x 12.3 human/mouse factor = 831 mg/kg mouse dose

We titrated up this dose to achieve a reproducible toxicity phenotype, ultimately landing on 1,500 mg/kg. This dose is in line with prior mouse studies of capecitabine pharmacodynamics using 1000-2000 mg/kg (*22, 75–77*). Consistent with our observations, the capecitabine product insert notes that doses of 1000 mg/kg in mice did not cause significant GI toxicity in pharmaceutical trials, while equivalent doses in humans result in significant GI side effects (*78*). Lower CAP toxicity in mice relative to humans may be due to metabolic differences (*21*), microbiome differences, or increased dihydropyrimidine dehydrogenase activity in intestinal tissue (*57*).

### CAP delivery in murine models

Buffer was prepared by dissolving 964 mg sodium citrate dihydrate and 139 mg of citric acid into 90 mL deionized water (DI), adjusting pH to 6.0 with HCl or NaOH, adding 5 g gum arabic, bringing final volume to 100 mL, and autoclaving for 20 min. CAP solution was prepared fresh daily (SPF experiments) or weekly (gnotobiotic experiments) by dissolving CAP into the buffer with continuous agitation and heating. Mice were given a single CAP bolus daily by oral gavage for all experiments except the CAP regional profiling experiment, where mice were gavaged twice daily. CAP was dosed by body weight, with CAP quantity determined such that 200 µL solution contained the CAP gavage dose (500, 1100 or 1500 mg/kg) for the largest mouse. Gavage needles were rinsed with DI in between each mouse, with different needles for each experimental group.

### CAP regional profiling experiment

11 mixed-sex mice were singly housed for one week, then gavaged with 500 mg/kg CAP (*n* = 6 total, 3 per sex) or 200 µL vehicle buffer (*n* = 5 total, 3 female, 2 male) twice daily at 7AM and 7PM (total CAP dosage of 1000 mg/kg/day) for two weeks. Mice were weighed daily, with no changes noted in vehicle vs CAP or over time (two-way ANOVA *p*>0.05). Baseline stool and endpoint stool, jejunum, ileum, cecum, proximal colon, and distal colon contents were collected, followed by DNA extraction and 16S rRNA gene sequencing.

### Establishment of CAP toxicity experiment

20 mice were gavaged with 1,500 mg/kg CAP (*n* = 12) or 200 µL buffer (*n* = 8) once daily. Half the mice were sacrificed at Day 4, with remaining mice sacrificed at Day 12. Endpoint small intestine, cecum, and colon contents were collected. 5 metrics were used to assess toxicity. (1) *Weight loss and survival time*. Mice were weighed daily and sacrificed at 15% weight loss (“survival time”) or at experimental endpoint, whichever came first. (2) *Anatomic measurements*. Mouse colon length, small intestine length, spleen weight, and gonadal fat pad weight were measured. (3) *Body composition*. Mouse body composition was measured at Days 0, 4, and 12 by EchoMRI with 1 primary accumulation. (4) *Lipocalin-2 ELISA*. Enzyme-linked immunosorbent assay was performed with Mouse Lipocalin-2/NGAL DuoSet ELISA kit using the manufacturer’s protocol with the following modifications a-c. (a) preweighed colon contents (50-100 mg) were combined with 1 mL PBS + 0.1% Tween 20 and vortexed for 20 min, then centrifuged for 10 min at 4°C and 12,000 rcf. For sample wells, 20 µL of extracted sample was added to 100 µL of reagent diluent and 6-fold serially diluted. (b) For standard wells, 125 pg/mL mouse lipocalin-2 standard with 2-fold serial dilutions was used, with two blank wells. (c) Absolute concentrations were calculated with linear fit to a log-log plot of mouse Lcn-2 concentration vs OD. (5) *Hand-foot syndrome*. Thermal hind paw hyperalgesia was measured as described in (*22*). Briefly, mice were placed on a 52°C hotplate and latency to rear paw lick, rear paw flick, or jump was recorded. *Sequencing*. We performed metagenomic sequencing of stool samples collected on Day 0 and Day 4. We focused on these samples to determine whether CAP-induced microbiota changes manifest by Day 4 and to maximize our statistical power to detect differences (20 stool samples were available at Day 4, whereas only 10 were available at Day 12 by design.)

### Broad-spectrum antibiotic CAP toxicity experiment

Antibiotic (AVNM) or Vehicle (Veh) water was prepared fresh weekly from autoclaved tap water supplemented with sucrose (0.5 g/L sucrose) ± antibiotics (1g/L ampicillin, 0.5g/L vancomycin, 1 g/L neomycin, 0.5 g/L metronidazole), followed by 0.2 μm filter sterilization. 24 mice were treated *ad libitum* with AVNM (*n* = 12) or Veh (*n* = 12) water for the duration of the experiment. After 1 week of antibiotics, mice were treated with 1,500 mg/kg CAP daily for 2 weeks. Weight was measured daily, and stool was taken at Days −7, 0, 3, 5 (relative to CAP start). Endpoint small intestine, cecum, and colon contents were collected. Endpoint cecal contents weight and gonadal fat pad weight were measured. To assess bacterial load, fecal slurries were generated by adding 100 mg Day 0 stool into 1 mL BHI+, vortexing for 1 min, and allowing fecal debris to settle for 5 min. 10 µL of supernatant was 10x serially diluted in BHI+, with 2.5 µL of each dilution spotted on BHI+ agar plates and incubated at 37°C for 5 days to quantify colony forming units (CFUs). All stool work was performed anaerobically.

### Germ-free vs conventional microbiota CAP toxicity experiment

To generate CONV-D mice, cecal contents from a single SPF donor was resuspended in 10% BHI+ aerobically and passed through a 100 μm filter, with 200 μL administered to GF mice followed by a 1-week engraftment period. CONV-R (*n* = 8), CONV-D (*n* = 8), and GF (*n* = 8) mice were treated with 1100 mg/kg CAP daily for 2 weeks. Weight was measured daily. Endpoint small intestine contents, cecal contents, colon contents, and large intestine length were collected for all mice, with endpoint small intestine length, spleen weight, cecal contents weight, and gonadal fat pad weight collected for CONV-D and GF mice.

### *preTA* CAP rescue experiments

Two independent experiments were conducted. In each, streptomycin water was prepared fresh weekly from autoclaved tap water supplemented with 5 g/L streptomycin, followed by 0.2 μm filter sterilization. 32 mice (Experient 1, **Fig. S11**) or 36 mice (Experiment 2, **Fig. 5**) were treated *ad libitum* with streptomycin water for the duration of the experiment. Overnight cultures of engineered *E. coli* strains (*E. coli* Δ*preTA*, *wt*, *preTA*^++^) were pelleted by centrifugation, washed with an equal volume of sterile 0.85% saline, pelleted by centrifugation, and resuspended in 1:10 sterile saline. Mice were gavaged with 200 µl bacterial suspension (Experiment 1: 18 Δ*preTA*, 18 *preTA*^++^; Experiment 2: 12 Δ*preTA*, 12 *wt*, 12 *preTA*^++^) by gavage following 1 day on strep water. One day post-colonization (“Day 0”), mice began 2 weeks of treatment with 1,500 mg/kg CAP daily. Weight was measured daily, and stool was taken at Days −1, 0, and at sacrifice. Endpoint small intestine, cecum, and colon contents were collected, along with colon length. Fecal slurries were generated by anaerobically adding 100 mg Day 0 stool into 1 mL LB + strep, vortexing 1 min, and allowing fecal debris to settle for 5 min. 10 µL of supernatant was 10-fold serially diluted in LB + strep, with 2.5 µL of each dilution spotted on EMB agar plates and incubated anaerobically at 37°C for 1 day to quantify *E. coli* CFUs.

### Gnotobiotic *preTA* CAP rescue experiment

Overnight cultures of engineered *E. coli* strains (*E. coli* Δ*preTA* and *E. coli preTA*^++^) were pelleted by centrifugation, washed with an equal volume of sterile 0.85% saline, pelleted by centrifugation again and resuspended in 1:10 sterile saline. Mice were gavaged with 200 µl bacterial suspension or saline only (*n* = 6 *E. coli* Δ*preTA*, *n* = 6 *E. coli preTA*^++^, *n* = 6 “mock”). One-week post-colonization, stool was collected and mice began 2 weeks of treatment with 1500 mg/kg CAP daily, with weight measured daily. Endpoint small intestine length and colon length were measured. Endpoint *E. coli* CFUs from Day 0 stool were quantified by culturing on MacConkey Agar as described previously (*16*). For group comparisons, Δ*preTA* and mock colonized mice were binned together into “mock/Δ*preTA*” (both contain no *preTA*).

### Capecitabine stool pharmacokinetics experiment

Streptomycin water was prepared from autoclaved tap water and supplemented with 5 g/L streptomycin, followed by 0.2 μm filter sterilization. 12 mice were treated *ad libitum* with streptomycin water for the duration of the experiment. Overnight cultures of engineered *E. coli* strains (*E. coli* Δ*preTA*, *wt*, *preTA*^++^) were pelleted by centrifugation, washed with an equal volume of sterile 0.85% saline, pelleted by centrifugation, and resuspended in 1:10 sterile saline. Mice were gavaged with 200 µl bacterial suspension (6 Δ*preTA*, 6 *preTA*^++^) by gavage following 1 day on strep water. One day post-colonization, baseline stool was collected, and mice were gavaged with 1,500 mg/kg CAP at 9AM. Stool was collected at 3, 5, and 7 hours post-gavage for CAP and 5-FU quantification.

### Gnotobiotic humanization CAP toxicity experiments

Two independent experiments were conducted. For each experiment, two donors with variable baseline *preTA* were selected to humanize 16 gnotobiotic mice (*n* = 8 per donor; 1 isolator per donor). Fecal slurries were generated anaerobically by adding 100 mg stool to 2 mL PBS, vortexing, and letting the mixture settle for 5 min. 1.5 mL of supernatant (“fecal slurry”) was transferred to a new tube on ice. 100 µL fecal slurry was orally gavaged into each mouse, with the researcher blinded to colonization group. One-week post-colonization, baseline stool was collected and mice began 2 weeks of treatment with 1,500 mg/kg CAP daily. Weight was measured daily. Endpoint stool, small intestine contents, cecal contents, colon contents, colon length, spleen weight, and gonadal fat pad weights were collected. Baseline stool was subjected to metagenomic sequencing to assess *preTA* status. For group comparisons, mice with no detected K17722/K17723 were called “*preTA* low”, with others called *“preTA* high.” Where stool was not evaluable (1/32 mice), mice were called as the *preTA* level of their cagemates.

### Statistical analysis

Statistical analysis was performed in R (v4.2.1) (*79*), with plots generated using ggplot2 (v3.5.1) and ggpubr (v0.6.0) (*66, 80*). All statistical tests are specified in the text or figure legends where used, with core tests summarized here. Linear mixed-effects models with day as a fixed effect and patient as a random effect to account for inter-subject baseline microbiome variability were used to identify statistically significant changes in alpha diversity, beta diversity, taxa, and genes with respect to time; the plotting unit ΔCLR/day thus represents the per-day change in CLR-transformed microbial or gene relative abundance. For differential mixed-effects analyses, samples prior to Day 0 (1^st^ day of treatment) were set to Day 0. Diversity, taxa, and genes were plotted with respect to time by LOESS regression with standard error shaded; patients where the taxa/gene of interest was not detected in any samples were excluded from these plots. PERMANOVA testing with treatment and patient as effects and patient as a strata was used to test for compositional differences in taxa and genes (CLR-Euclidean ordination). Comparisons between two groups were performed with Mann-Whitney U test or ANOVA. Comparisons between three groups were performed with Kruskal-Wallis testing. Two-way ANOVA was used to identify significant group differences in mouse weight with respect to time. Mouse weights are plotted as mean with standard error bars; days after the first mouse sacrifice due to 15% weight loss are excluded from these plots. Spearman correlation was used to identify significant relationships between continuous variables. Survival distributions were compared with the Mantel-Cox test. pROC (v1.18.5) command roc was for calculating sensitivity, specificity, and AUROC (*81*). Significance was determined as *p*-value < 0.05 (individual tests) or Benjamini-Hochberg FDR < 0.2 (multiple hypothesis correction of taxa/genes).

## List of Supplementary Materials

Figs. S1-S17

Tables S1-S3

Data Files S1-S8

## Acknowledgements

Gnotobiotic experiments were performed at the UCSF Gnotobiotics Core Facility. Sequencing was performed at Chan-Zuckerberg Biohub-San Francisco and the UCSF Center for Advanced Technology. Diagrams were created with BioRender.com.

## Funding

National Institutes of Health Grant F30CA257378 (TSK)

National Institutes of Health Grants R01HL122593, R01CA255116, R01DK114034 (PJT)

Parkinson’s Disease Foundation Grant PF-PRF-932695 (CAO)

## Author contributions

Conceptualization: KRT, WAK, TSK, CEA, EVB, PJT

Methodology: KRT, TSK, CN, CAO, TH, EFO, VU, JAT, DD, PJT

Investigation: KRT, TSK, CAO, EFO, TH, MDS, PS, DS

Visualization: KRT

Funding acquisition: WAK, AV, CEA, PJT

Project administration: WAK, PJT

Supervision: CAO, CN, VU, CEA, PJT

Writing – original draft: KRT, PJT

Writing – review & editing: All authors

## Competing interests

W.A.K. has received research funding (institution) from Pfizer; there is no direct overlap with the current study. P.J.T. is on the scientific advisory boards of Pendulum, Seed and SNIPRbiome; there is no direct overlap between the current study and these consulting duties. C.E.A served on the scientific advisory board of Pionyr Immunotherapeutics and has received research funding (institution) from Bristol Meyer Squibb, Erasca, Gossamer Bio, Guardant Health, Kura Oncology, Merck and Novartis; there is no direct overlap with the current study. E.V.B. is on the medical advisory board for Fight CRC; there is no direct overlap with the current study. All other authors declare no competing interests.

## Data and materials availability

This study did not generate new reagents. All code and data for generating figures is on GitHub at https://github.com/turnbaughlab/2024_Trepka_GO. The accession number for the sequencing data reported in this paper is NCBI Sequence Read Archive BioProject PRJNA1169175.

## Supplementary Materials

**Figure S1:**
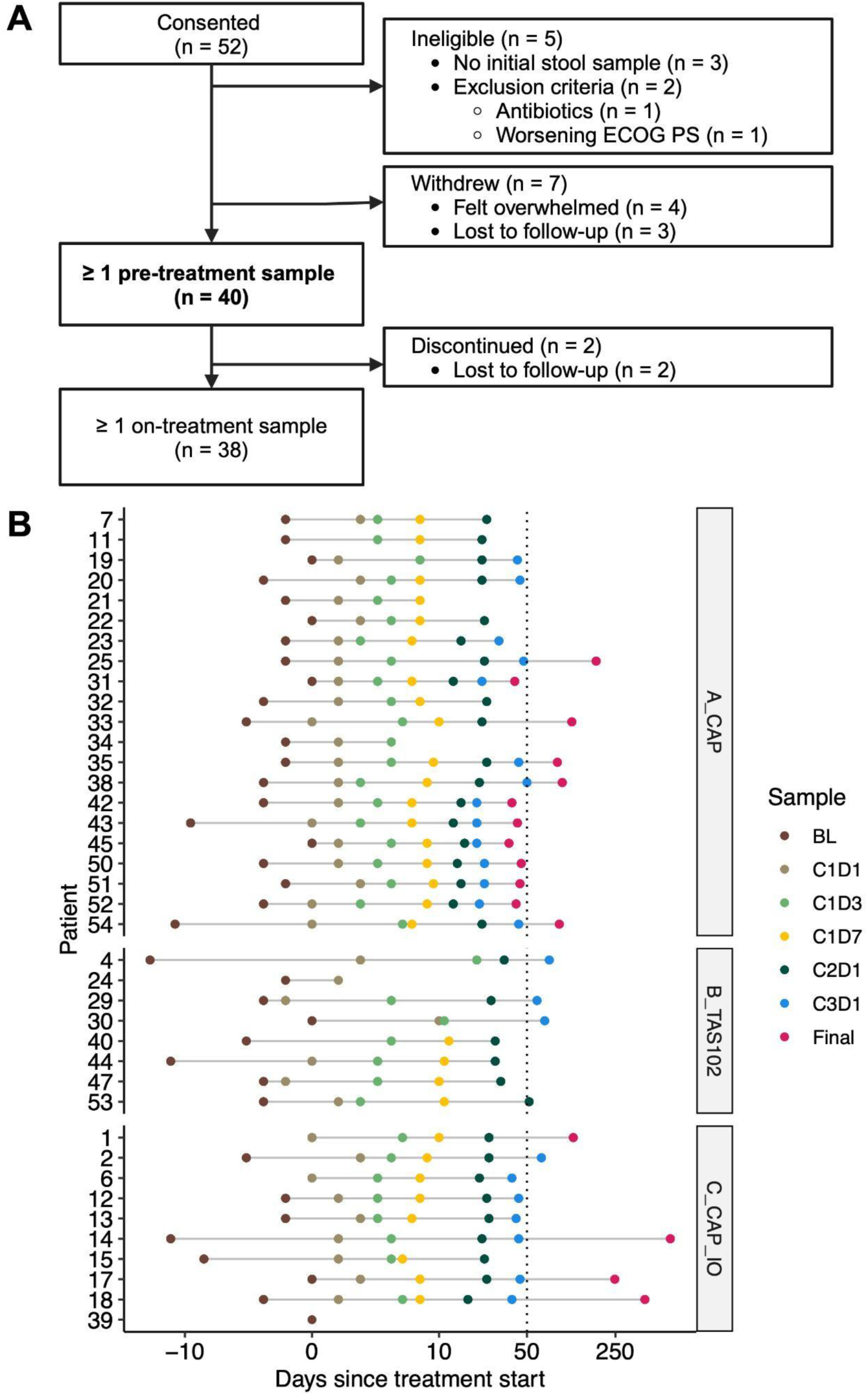
GO study design. **(A)** Flow diagram of patients consented to GO study. **(B)** Day of evaluable stool samples from each patient relative to Day 0, the first day of Cycle 1 of chemotherapy. Each dot represents a single stool sample, and each color represents a study-defined timepoint. Samples to the right of the vertical dotted line at Day 50 were excluded from mixed-effects modeling. The x axis is pseudo-log transformed.

**Figure S2:**
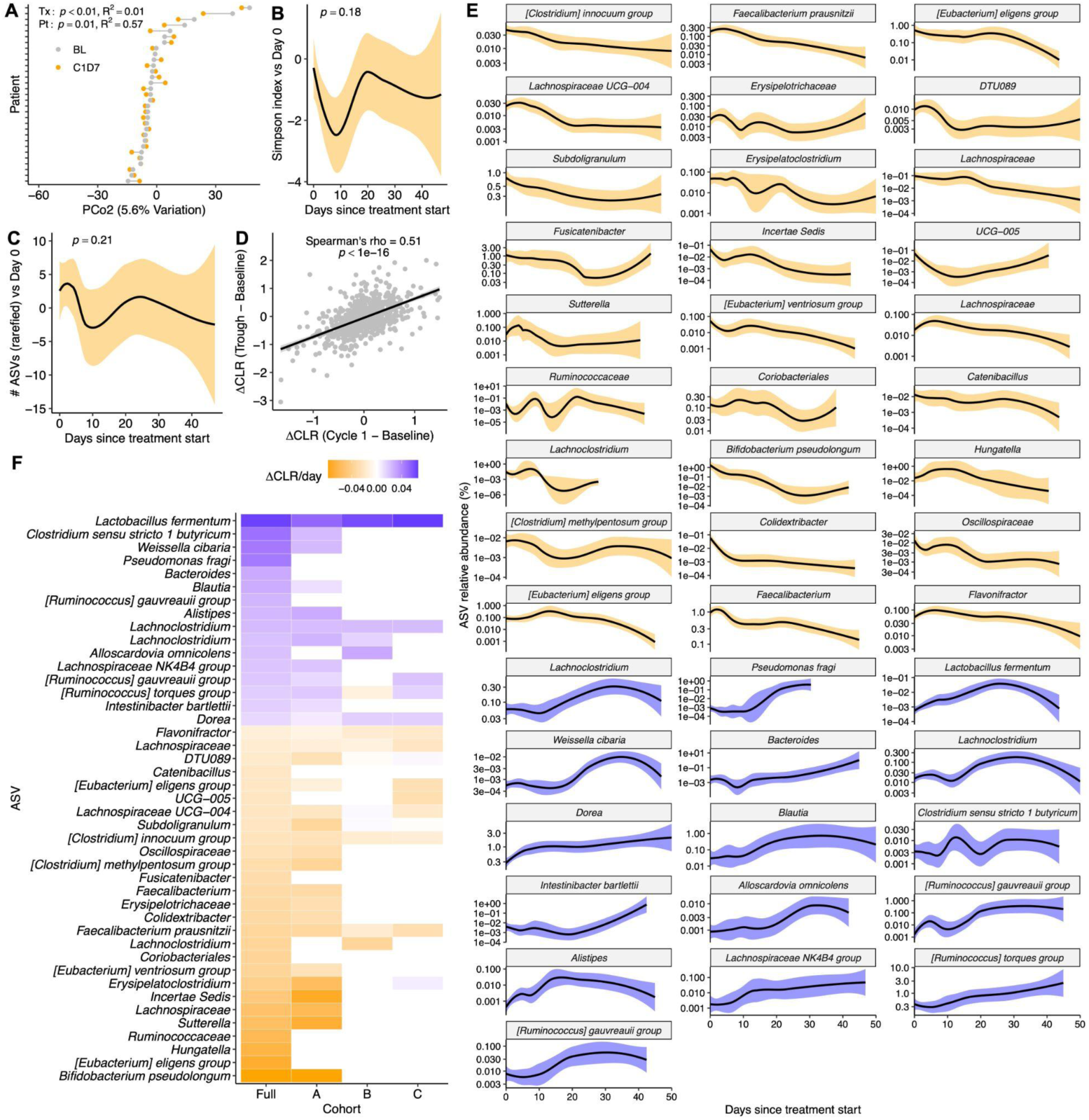
Oral fluoropyrimidine treatment impacts gut microbiome composition in GO study patients. **(A)** Reproducible shifts in gut microbial amplicon sequence variants (ASV) are observed across study participants between BL and C1D7 samples on the second principal coordinate (PCo2) of central log ratio (CLR)-transformed Euclidean distances. *p*-value: PERMANOVA with Patient ID as stratum, all samples. **(B,C)** Alpha diversity metrics inverse Simpson index **(B)** and number of ASVs **(C)** during the study, normalized to baseline. *p*-values: mixed-effects model, Diversity ∼ Day + 1|Patient. **(D)** Correlation between taxa changes observed in Cycle 1 and Trough relative to Baseline. *p-*value: Spearman’s rank correlation. **(E)** Time course of all significantly altered ASVs from Fig. 1C, with ASVs that increased/decreased during treatment in blue/orange, respectively. Solid lines with shading represent LOESS interpolation mean±sem. **(F)** Heatmap of differential ASV abundance in the full cohort and each sub-cohort. Displayed ASVs are those that are significantly altered in the full cohort (Fig. 1C). Shading color indicates treatment effect (orange, depleted with treatment; blue, enriched), with white indicating the ASV was detected in fewer than 3 patients in the sub-cohort.

**Figure S3:**
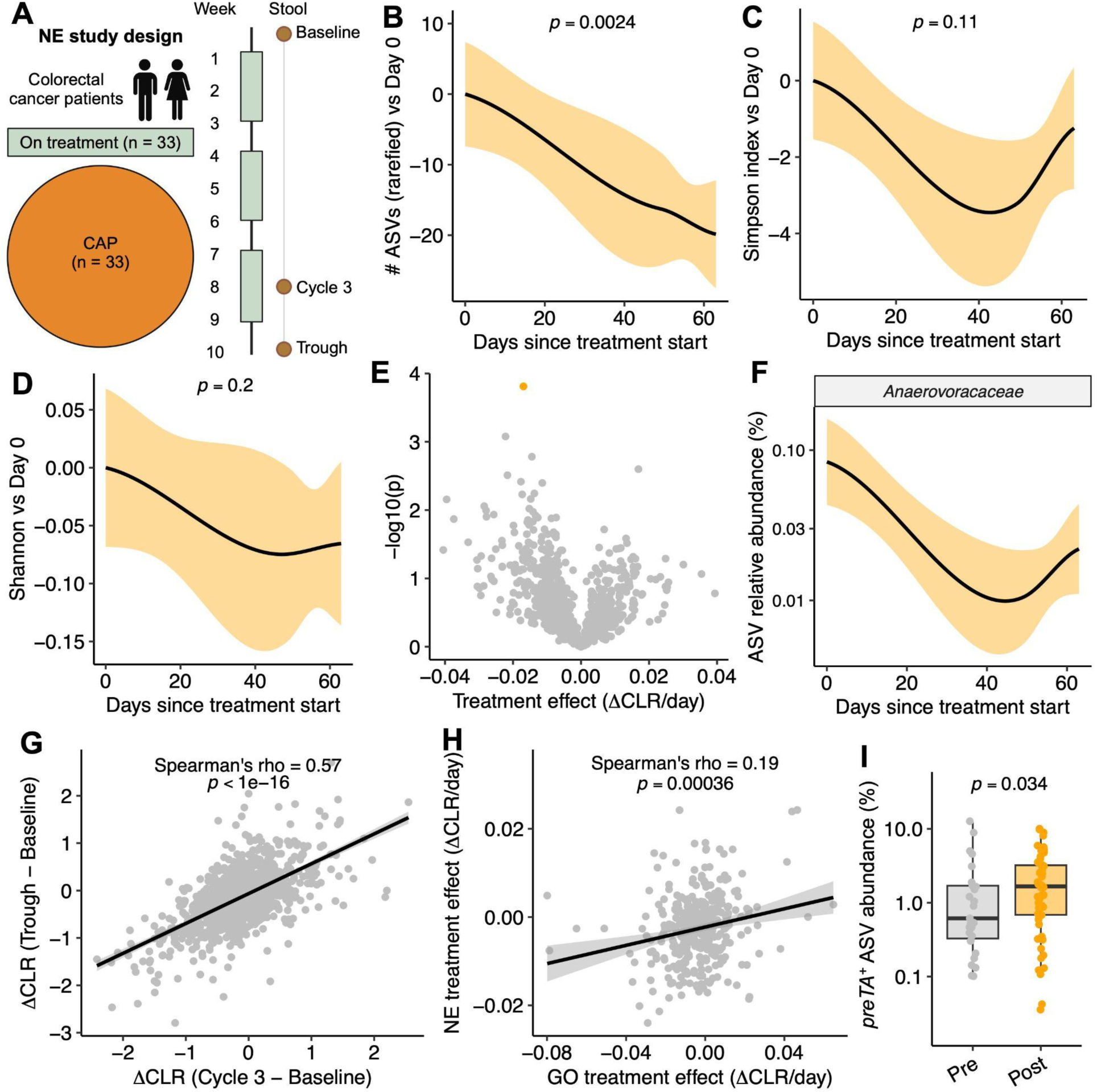
Oral fluoropyrimidine treatment impacts gut microbiome composition in a European cohort of colorectal cancer patients. **(A)** Netherlands (NE) study design depicting patients treated with capecitabine (CAP). Created with BioRender.com. **(B-D)** Change in alpha diversity metrics number of rarefied ASVs **(B)**, inverse Simpson index **(C),** Shannon index **(D)** during the study, normalized to baseline. *p*-values: mixed-effects model, Diversity ∼ Day + 1|Patient. **(E)** Volcano plot of microbial amplicon sequence variants (ASVs) with respect to treatment time (mixed-effects model, central log ratio (CLR)-transformed Abundance ∼ Day + 1|Patient). Orange represents the only significantly depleted ASV after treatment (false discovery rate (FDR) < 0.2). **(F)** Time course of significantly altered *Anaerovoracaceae* ASV. **(G)** Correlation between taxa changes observed in Cycle 3 and Trough relative to Baseline. **(H)** Correlation between taxa changes observed in GO and NE studies. **(I)** Sum of *preTA*^+^ *Escherichia*, *Anaerostipes*, *Eubacterium, Citrobacter* ASV relative abundance. *p*-value: one-sided Mann-Whitney U test. **(B-D,F)**: Solid lines with shading represent LOESS interpolation mean±sem. **(G,H)**: Solid lines with shading represent a linear regression best fit with 95% confidence interval, with Spearman correlation *p*-values.

**Figure S4:**
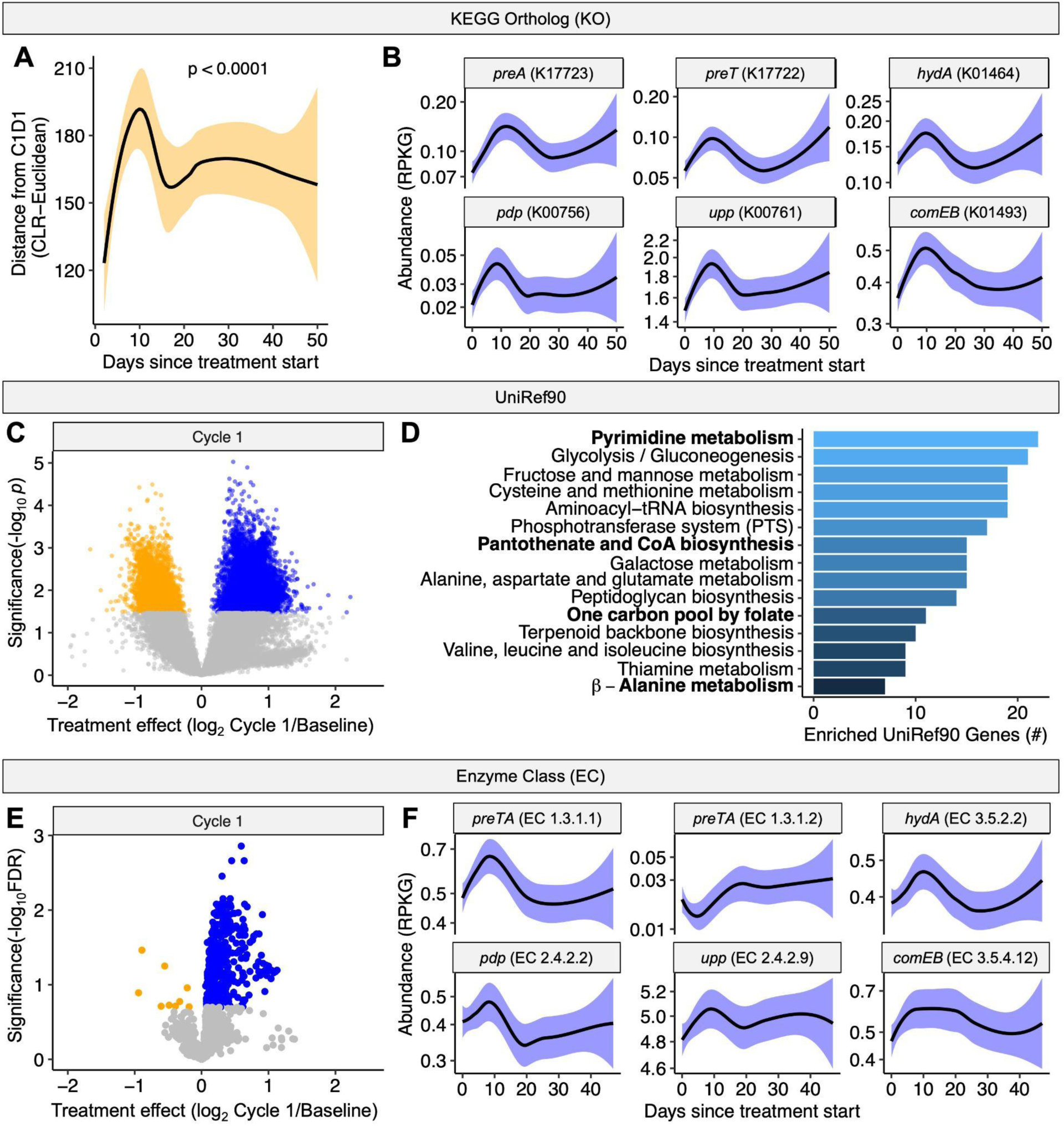
Oral fluoropyrimidine treatment selects for pyrimidine metabolism genes. **(A)** Microbial community distance to Cycle 1 Day 1 location calculated per-patient (central log ratio-Euclidean ordination). *p*-value: mixed-effects model, Distance ∼ Day + 1|Patient. **(B)** Time course of significantly altered KEGG Ortholog gene families from pyrimidine metabolism, pantothenate and CoA biosynthesis, and β-alanine metabolism pathways. **(C)** Volcano plot of UniRef90 gene families detected in at least 50% of samples with respect to treatment time. Points represent significantly enriched (blue) and depleted (orange) gene families (FDR < 0.2). **(D)** Gene set enrichment analysis of significantly enriched UniRef90 gene families from **(C)**. UniRef90 gene sets with *p* < 0.1 are displayed. Fluoropyrimidine metabolism-related pathways are bolded. **(E)** Volcano plot of Enzyme Classes (EC) detected in at least 50% of samples with respect to treatment time. Points represent significantly enriched (blue) and depleted (orange) ECs (FDR < 0.2). **(F)** Time course of ECs corresponding to genes from **(B)**. **(A,B,F)** Solid line with shading represents LOESS interpolation mean±sem.

**Figure S5:**
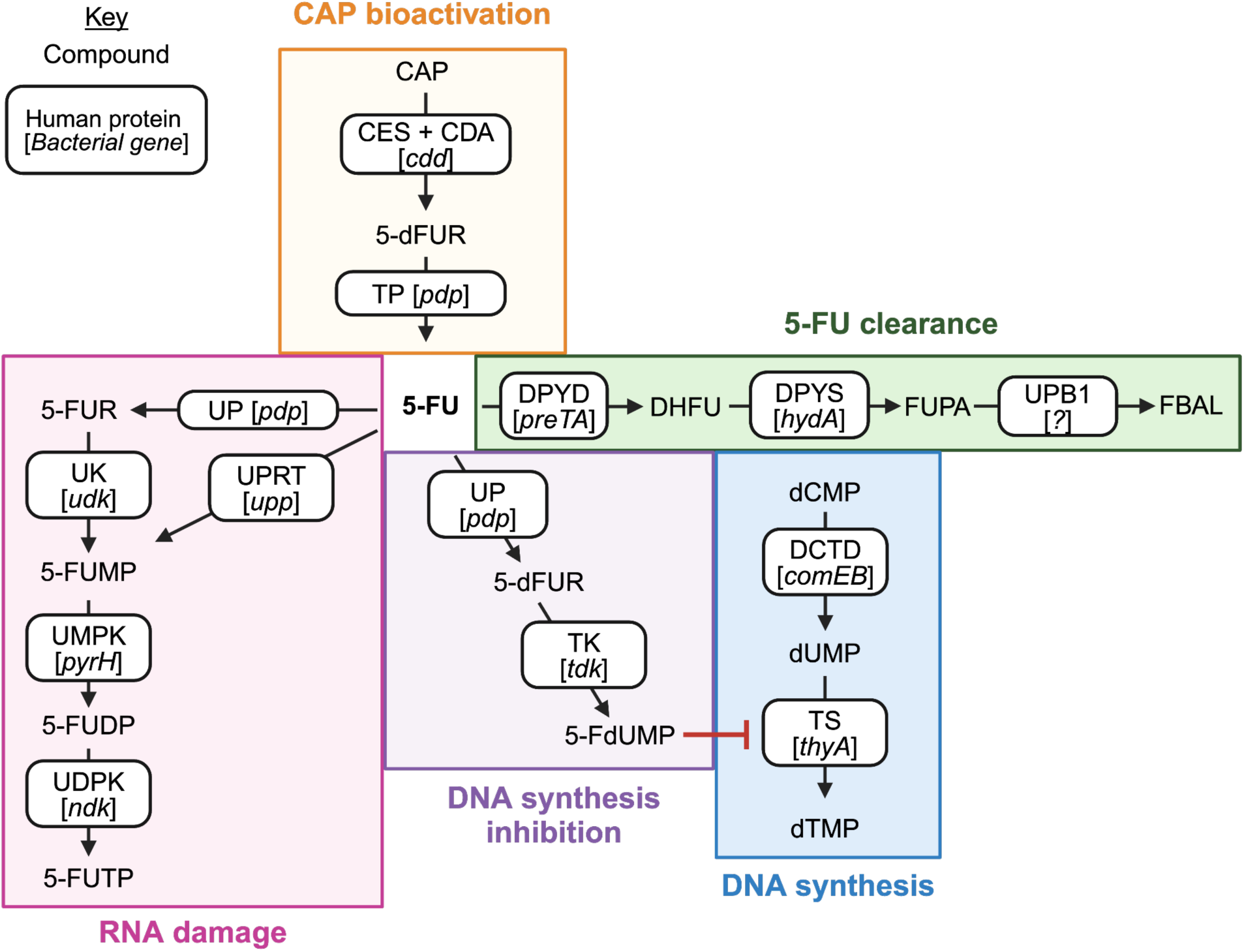
Fluoropyrimidine bioactivation and metabolism. Created with BioRender.com. Text in boxes represents enzymes responsible for a transformation, formatted as human protein [*bacterial gene*]. Text outside boxes represents fluoropyrimidine metabolites. Arrows indicate directionality to produce terminal metabolites responsible for cytotoxicity or clearance, but do not indicate that a reaction is irreversible. **CAP bioactivation**. In humans, capecitabine (CAP) is de-esterified into 5-DFCR by liver carboxyesterase (CES), followed by deamination into 5-dFUR by liver and tumor cytidine deaminase (CDA) (*1*). In bacteria, *Bacillus subtilis cdd* encodes a tetrameric cytidine deaminase that directly converts CAP to 5-dFUR, a capacity also shared by other enzymes cloned from gut metagenomic sequences (*2*). In humans, thymidine phosphorylase (TP) converts 5-dFUR to 5-FU (*1*); in bacteria, pyrimidine nucleoside phosphorylase genes (*pdp*, including *upp* in *E. coli*) encode the responsible enzymes (*3, 4*). **5-FU clearance**. In humans and bacteria, 5-FU is cleared to DHFU by homologous dihydropyrimidine dehydrogenases (DPYD, encoded by *preTA* in bacteria including *E. coli* and *Anaerostipes hadrus*) (*5*). DHFU is further converted to FUPA by homologous dihydropyrimidases (DPYS, encoded by *A. hadrus hydA*) (*6*). In humans, FUPA is then converted to fluoro-beta-alanine by beta-ureidopropionase (UPB1) prior to renal clearance (*7*). **DNA synthesis**. *De novo* DNA synthesis requires conversion of dUMP to dTMP by thymidylate synthase (TS, encoded by *E. coli thyA*) (*8*). Upstream, dCMP stores can be converted to dUMP by dCMP deaminase (DCTD, encoded by *B. subtilis comEB*) (*9*). **DNA synthesis inhibition**. DNA synthesis inhibition is a primary mechanism-of-action of 5-FU (*10*). In humans and bacteria, 5-FU is converted to 5-dFUR by uridine phosphorylase (UP, encoded by bacterial *pdp* including *E. coli udp*), which is then converted to 5-FdUMP by thymidine kinase (TK, encoded by *E. coli tdk*) (*11*). 5-FdUMP then forms an inhibitory ternary complex with thymidylate synthase (TS, encoded by *E. coli thyA*), blocking DNA synthesis (*8*). **RNA damage**. In humans and bacteria, 5-FU is converted directly to 5-FUMP by UPRT (UPRT, encoded by *E. coli* and *Mycobacterium spp. upp*) (*5, 12*). Alternatively, 5-FU can be converted to 5-FUR by uridine phosphorylase (UP, encoded by bacterial *pdp* including *E. coli udp*), which is then converted to 5-FUMP by uridine kinase (UK, encoded by *E. coli udk*) (*11*). 5-FUMP is sequentially phosphorylated to 5-FUDP and 5-FUTP by uridine monophosphate kinase (UMPK, encoded by *E. coli pyrH*) and uridine diphosphate kinase (UDPK, encoded by *E. coli ndk*) (*11*). 5-FUTP disrupts RNA processing, resulting in cytotoxicity (*13*).

**Figure S6:**
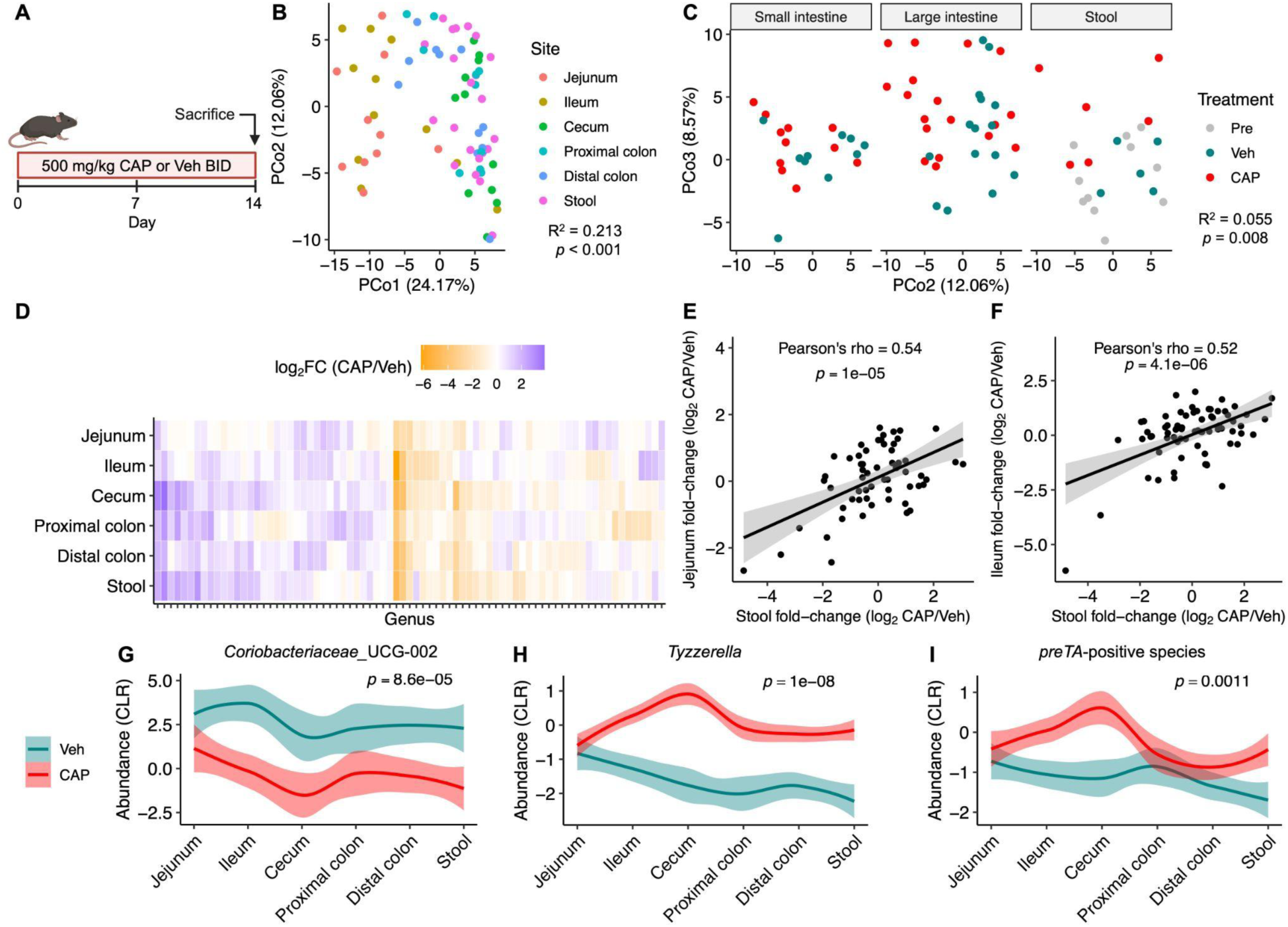
CAP alters murine gut microbiota composition along the gastrointestinal (GI) tract. **(A)** CAP GI mouse model (created with BioRender.com). Mixed-sex mice were gavaged daily with 500 mg/kg CAP (*n* = 6) or Vehicle (Veh, *n* = 5) twice daily (BID), for a total of 1000 mg/kg CAP/day. Stool was collected at Day 0, and all gastrointestinal contents (jejunum, ileum, cecum, proximal colon, distal colon, stool) were collected at Day 14. **(B)** Microbial genera compositions separate by body site using principal coordinate analysis (PCoA) with central log ratio (CLR)-transformed Euclidean distances. p-value: PERMANOVA with Mouse ID as stratum. **(C)** Microbial genera compositions separate by treatment per body site (Small intestine: Jejunum, Ileum; Large intestine: Cecum, Proximal colon, Distal colon; Stool) using PCoA with CLR-Euclidean distances. *p*-value: PERMANOVA of all data combined with Mouse ID as stratum. **(D)** Heatmap of endpoint differential genera abundance for each body site, with genera ordered by hierarchical clustering (Ward D2). **(E-F)** Correlation between taxa changes observed in jejunum (**E**) or ileum (**F**) vs stool. p-value: Pearson’s correlation. **(G-I)** Abundance of most depleted genus in stool (**G**), most enriched genus in stool (**H**), and *preTA*-positive species across body sites (**I**). *p*-values: mixed-effects model, Abundance ∼ Treatment + 1|Tissue. Solid line with shading represents LOESS interpolation mean±sem.

**Figure S7:**
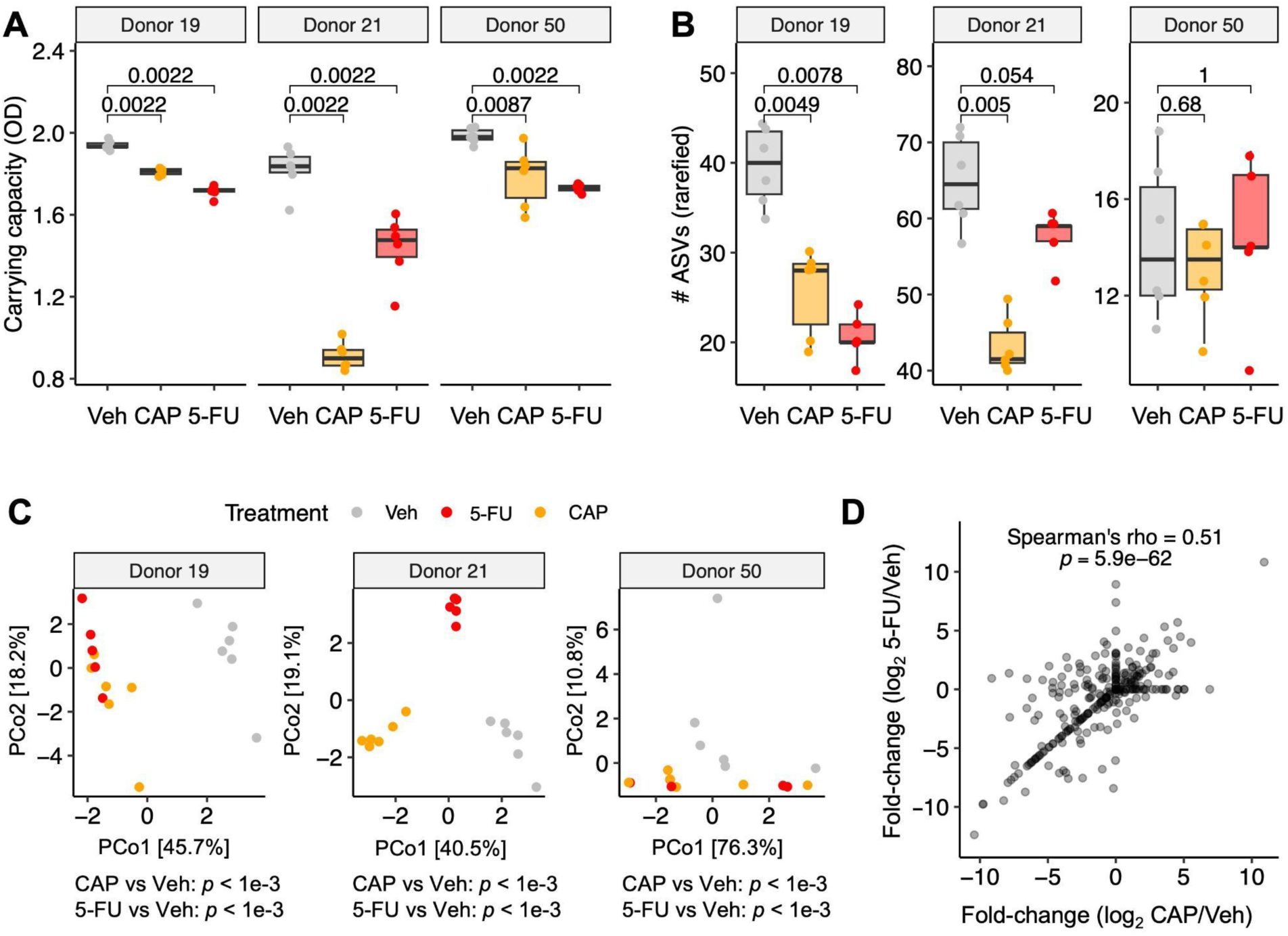
Fluoropyrimidines alter human gut microbiota composition *ex vivo*. **(A)** Carrying capacity (optical density, OD) and **(B)** alpha diversity (number of amplicon sequence variants, ASVs) of stool-derived *ex vivo* communities incubated with Vehicle (Veh), CAP, or 5-FU. *p*-value: Mann-Whitney U test. **(C)** Principal coordinate plots of ASV abundances for *ex vivo* communities incubated with Veh, 5-FU, or CAP (central log ratio (CLR)-Euclidean ordination). *p*-value: Treatment PERMANOVA relative to Vehicle. **(D)** CAP- and 5-FU-induced changes in amplicon sequence variant abundance are highly correlated. *p*-value: Spearman correlation. Each point represents a single *ex vivo* community **(A-C)** or a single amplicon sequence variant across communities from a given donor **(D)**. To calculate log_2_ fold change, a pseudocount of 0.00001 abundance was added to the numerator and denominator.

**Figure S8:**
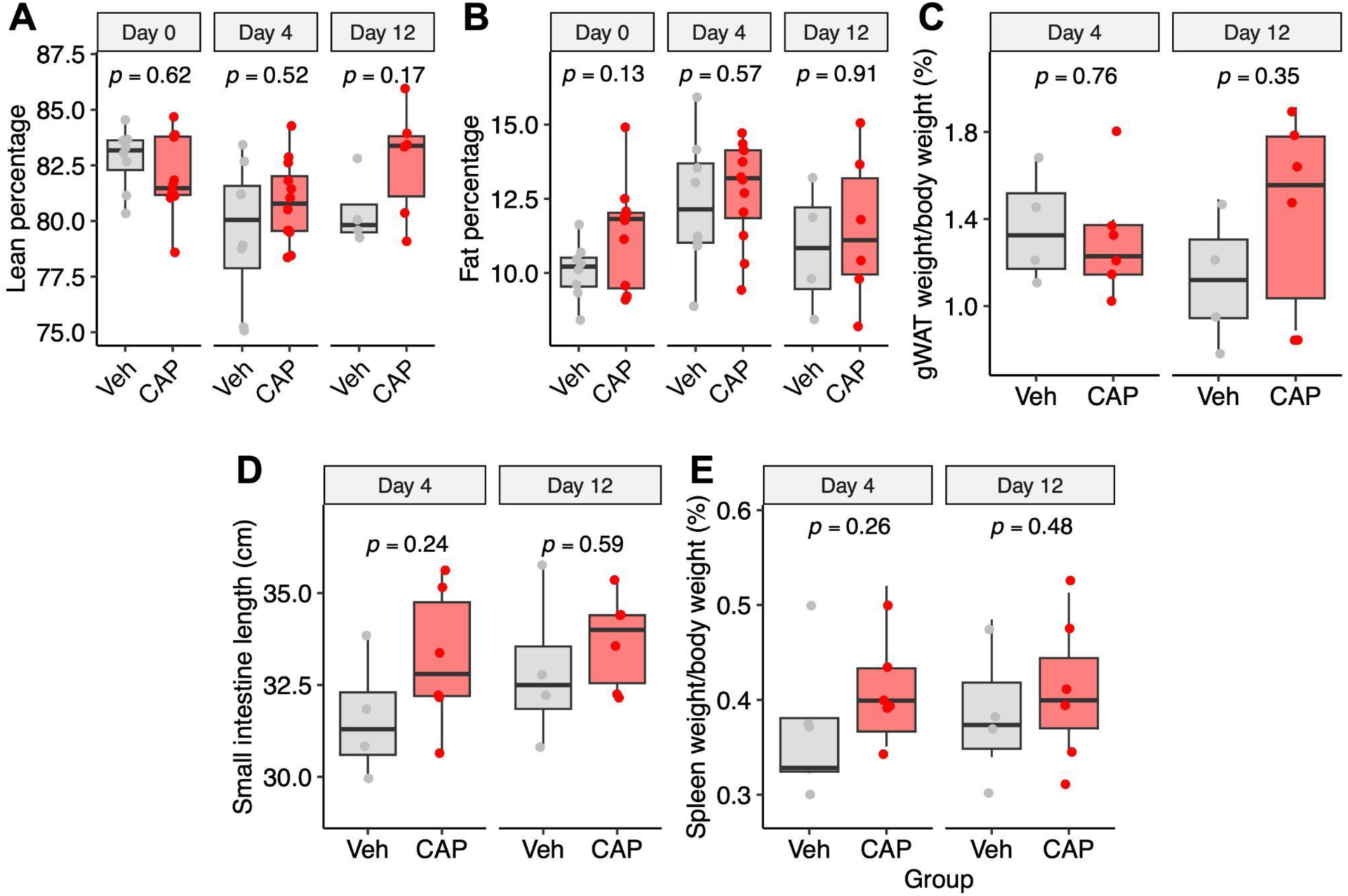
A mouse model of capecitabine (CAP) toxicity. **(A-B)** Body composition by EchoMRI at Day 0, Day 4 and Day 12 of CAP treatment: lean percentage **(A)** and fat percentage **(B)**. Toxicity endpoints at Day 4 and Day 12 of CAP treatment: gonadal white adipose tissue (gWAT) weight/body weight **(C)**, small intestine length **(D)**, and spleen weight/body weight **(E)**. *p-*values: Mann-Whitney U test, CAP vs Veh.

**Figure S9:**
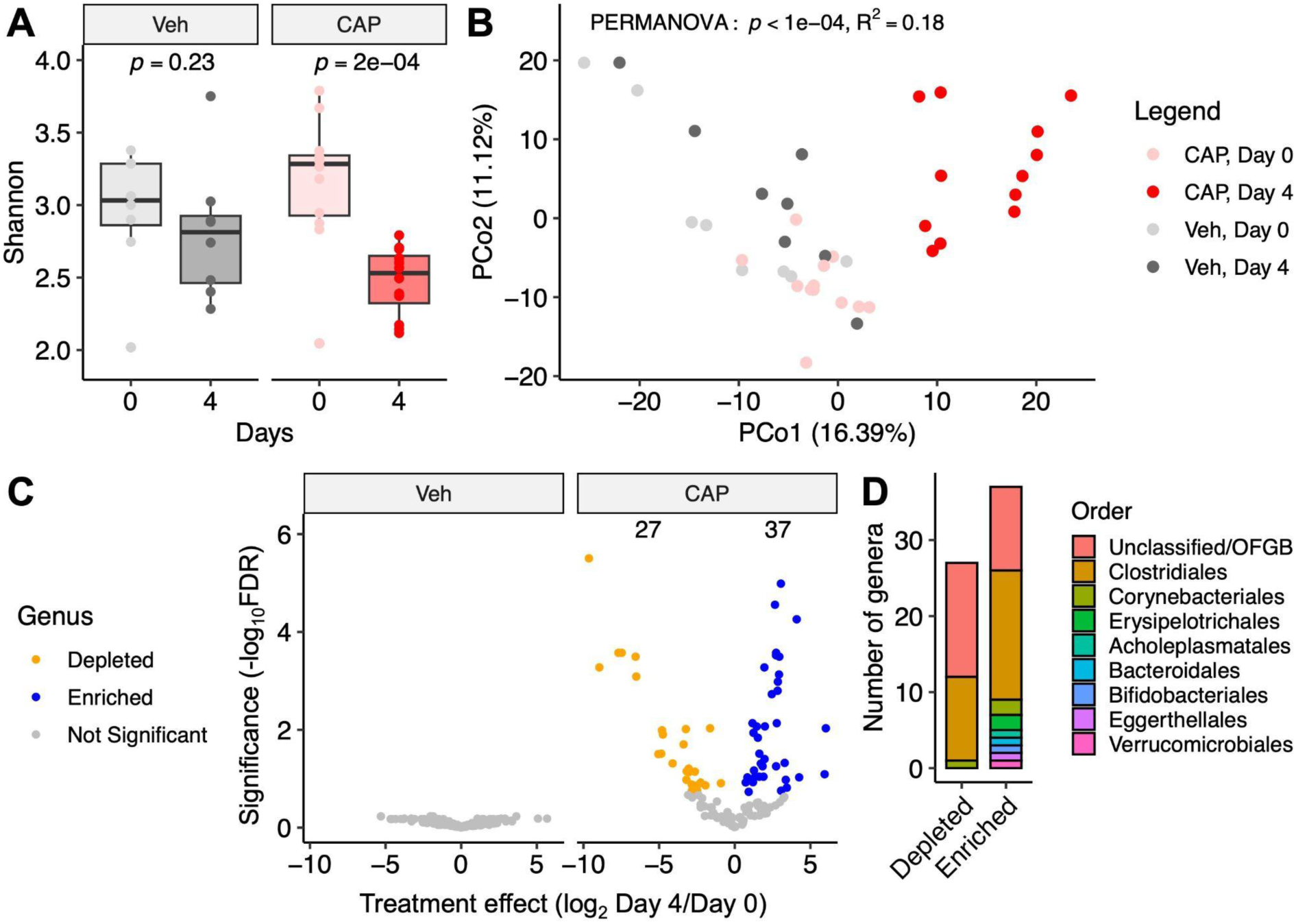
Capecitabine (CAP) alters murine gut microbiota composition in a toxicity model. **(A)** Shannon index during the experiment. *p*-value: Mann-Whitney U test. **(B)** Shifts in gut microbial genera are observed between Day 0 and 4 in CAP-treated mice using principal coordinate analysis (PCoA) of central log ratio (CLR)-transformed Euclidean distances. *p*-value: PERMANOVA with Mouse ID as stratum. **(C)** Volcano plot of microbial genera with respect to treatment time, comparing Day 0 and 4. *p*-value: mixed-effects model, CLR-transformed Abundance ∼ Day + 1|Mouse. Points represent enriched (blue) and depleted (orange) genera after treatment (false discovery rate (FDR) < 0.2). **(D)** Order membership of enriched and depleted genera from **(C**).

**Figure S10:**
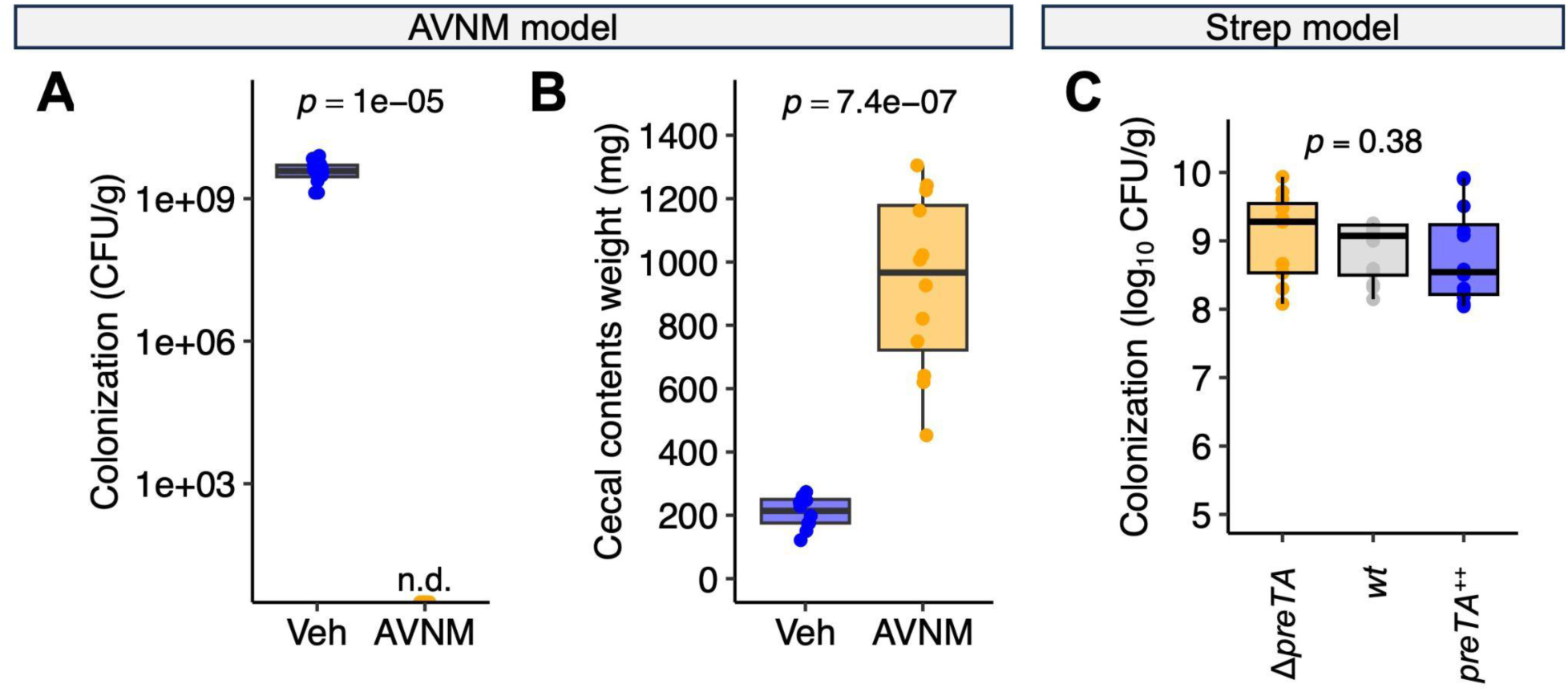
Microbial load in antibiotic depletion models. (**A-B**) An ampicillin, vancomycin, neomycin, and metronidazole (AVNM) cocktail depletes the gut microbiota, measured using stool colonization at Day 0 (1 week post-AVNM) **(A)** and cecal contents weight at sacrifice **(B)**. **(C)** *E. coli ΔpreTA*, *wt*, and *preTA*^++^ similarly colonize mice treated with 5 g/L streptomycin. *p-*values: Mann-Whitney U test **(A,B)**, Kruskal-Wallis test (**C**).

**Figure S11:**
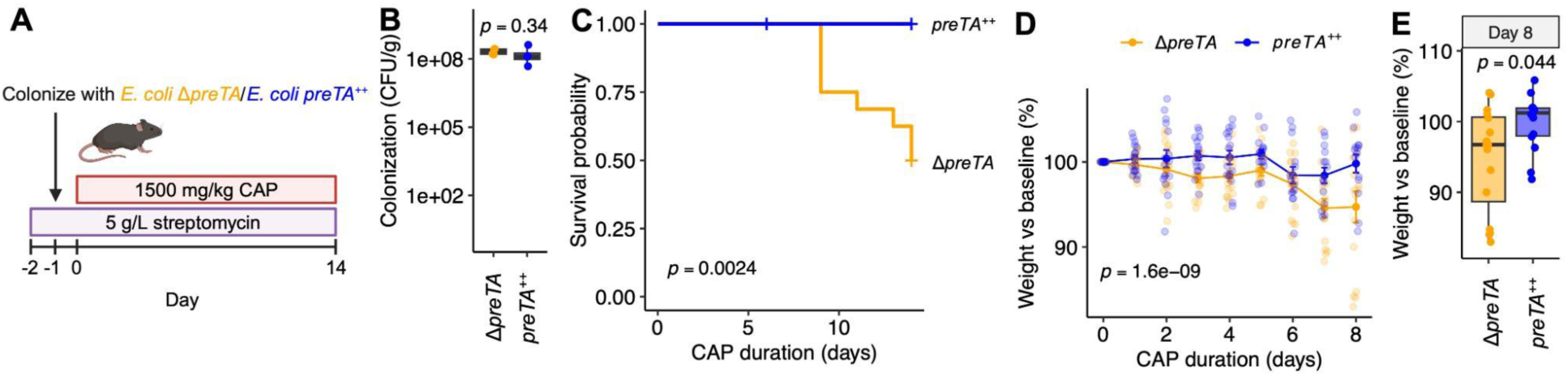
Microbial *preTA* rescues capecitabine (CAP) toxicity in a streptomycin-treated mouse model. **(A)** *preTA* streptomycin (strep) colonization CAP toxicity model in female-only C57BL/6J mice (separate experiment from Fig. 5; created with BioRender.com). Female mice were treated with streptomycin for 1 day, gavaged with *E. coli* Δ*preTA* (*n* = 18) or *preTA*^++^ (*n* = 18), then gavaged daily with 1500 mg/kg CAP. **(B)** *E. coli* colonization level by cage. **(C)** Survival curve. **(D-E)** CAP-induced weight loss **(D)** and weight on Day 8 **(E**). *p*-values: Mann-Whitney U test **(B,E)**, Mantel-Cox test **(C)**, group term from two-way ANOVA (**D**).

**Figure S12:**
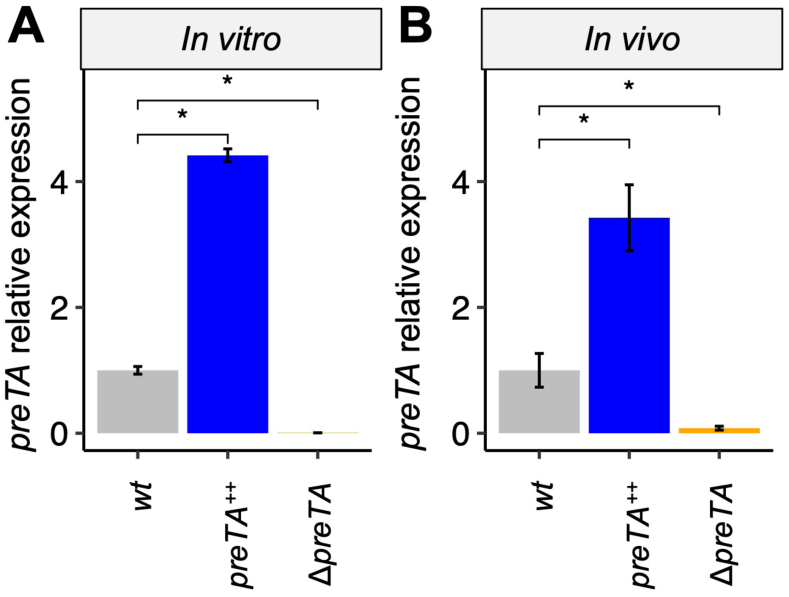
*preTA^++^ E. coli* transcript levels are significantly higher than *wt E. coli*. *preTA* expression levels in pure culture **(A)** and mice treated with 5 g/L streptomycin followed by colonization **(B)**. All expression values were normalized to housekeeping gene *rrsA*, then divided by mean *wt preTA* expression. Values are mean±sem, \**p*<0.05, Mann-Whitney U test.

**Figure S13:**
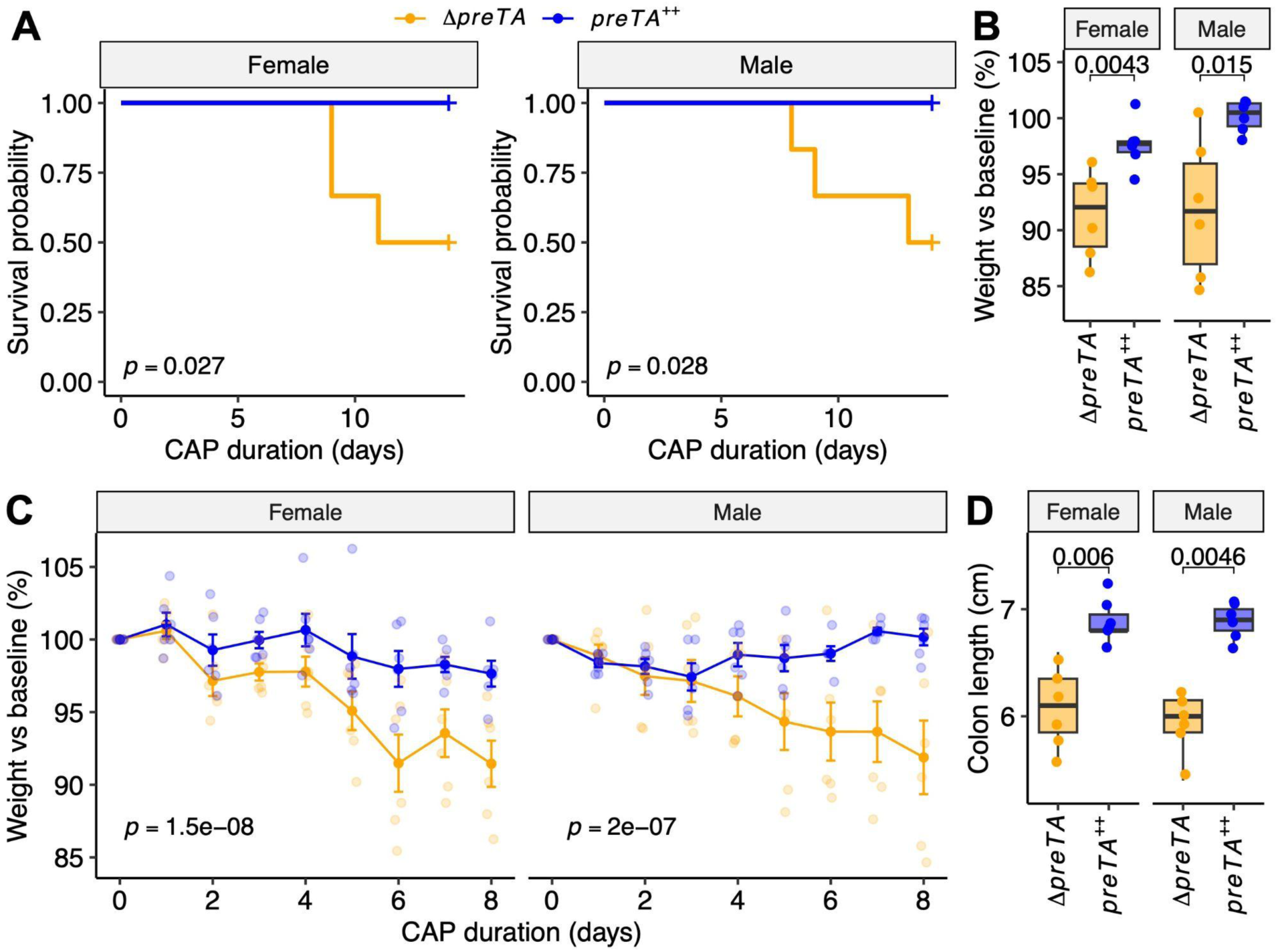
Bacterial *preTA* rescues capecitabine (CAP) toxicity in both male and female mice. Mixed-sex mice were treated with streptomycin for 1 day, gavaged with *E. coli* Δ*preTA* (*n* = 12) or *preTA*^++^ (*n* = 12), then gavaged daily with 1500 mg/kg CAP (same data as **Figs. 5A-E**). **(A)** CAP survival curves. **(B)** Endpoint colon lengths. **(C)** CAP-induced weight loss. **(D)** Mouse weight on Day 8. *p*-values: one-sided Mantel-Cox test **(A)**, Mann-Whitney U test **(B,D)**, group term from two-way ANOVA (**C**).

**Figure S14:**
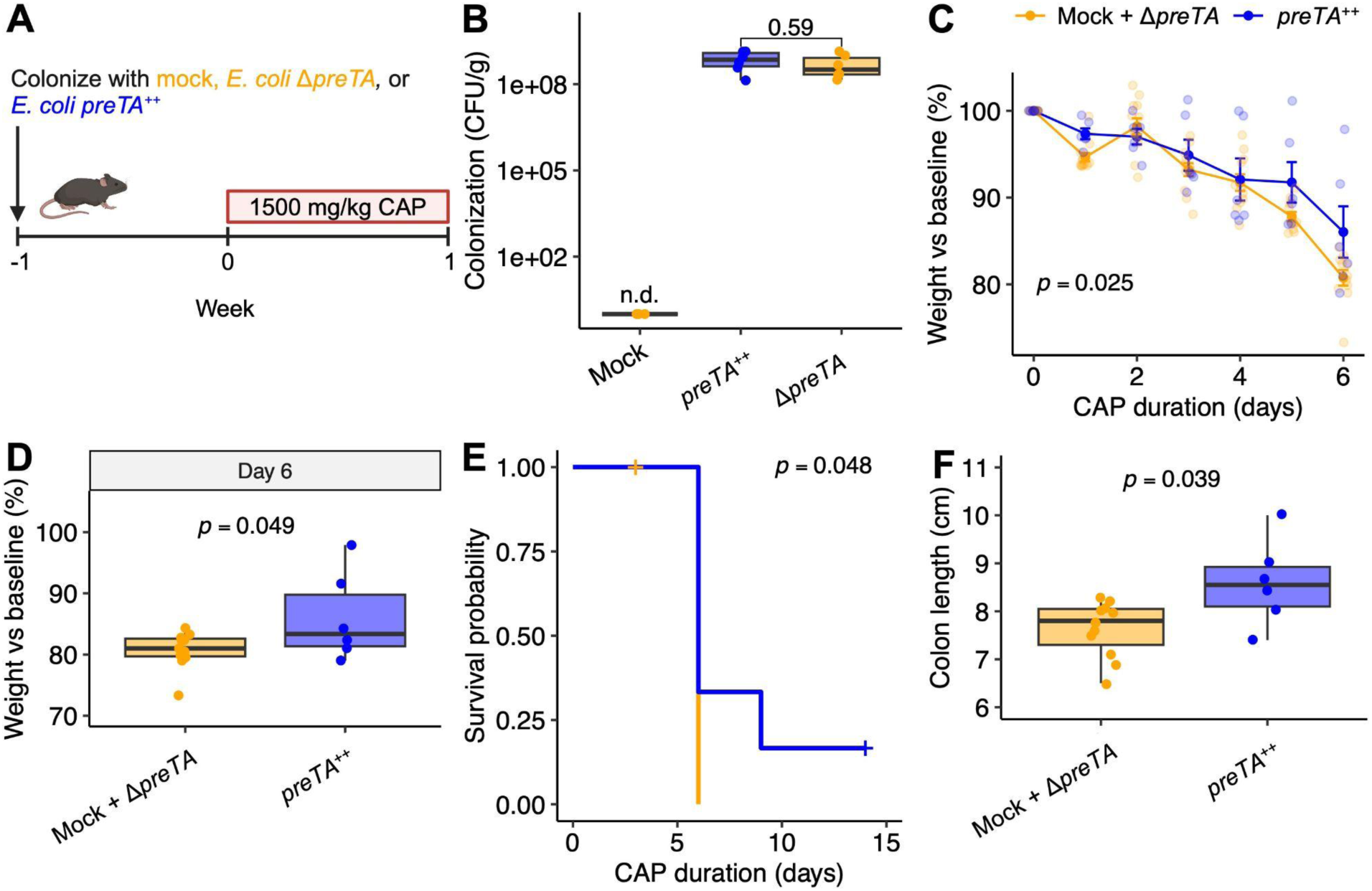
Microbial *preTA* rescues capecitabine (CAP) toxicity in a gnotobiotic mouse model. **(A)** Experimental design of gnotobiotic mice colonized with mock (*n* = 6), Δ*preTA E. coli* (n = 6), or *preTA^++^ E. coli* (n = 6) for 1 week prior to treatment with 1500 mg/kg CAP daily by oral gavage. Created with BioRender.com. **(B)** *E. coli* stool colonization level at baseline. n.d. = not detected. **(C)** CAP-induced weight loss. *p-*value: two-way ANOVA. **(D)** Mouse weight on Day 6. *p*-value: ANOVA **(E)** Survival curve. *p*-value: Mantel-Cox test. **(F)** Colon length at sacrifice. **(B,F)**: *p*-value: Mann-Whitney U Test.

**Figure S15:**
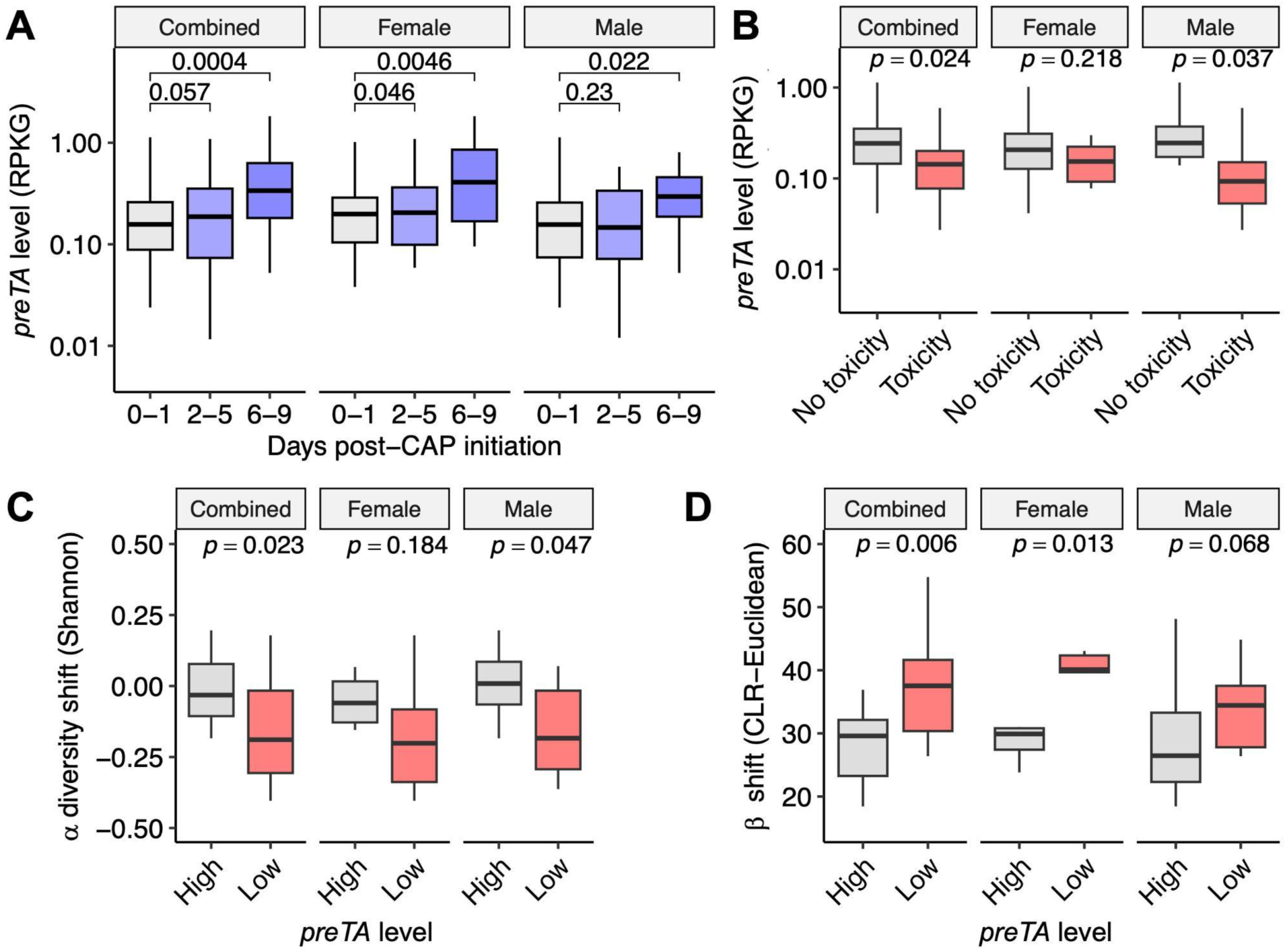
Capecitabine-*preTA* interactions are detectable for each sex. **(A)** *preTA* levels during Cycle 1 of CAP treatment. *p*-value: one-sided paired Student’s *t*-test. **(B)** Patients experiencing toxicity have lower baseline stool bacterial *preTA* levels (*n* = 40 combined, 20 male, 20 female). *p*-value: one-sided Mann-Whitney U test. **(C)** *preTA* level (above/below mean) vs change in alpha diversity during Cycle 1 (C7D1-C1D1 Shannon index, *n* = 30 combined, 18 male, 12 female). **(D)** *preTA* level (above/below mean) vs change in beta diversity during Cycle 1 (C7D1-C1D1 CLR-Euclidean distance, *n* = 30 combined, 18 male, 12 female). **(C,D)** *p*-value: one-sided Wilcoxon signed-rank test.

**Figure S16:**
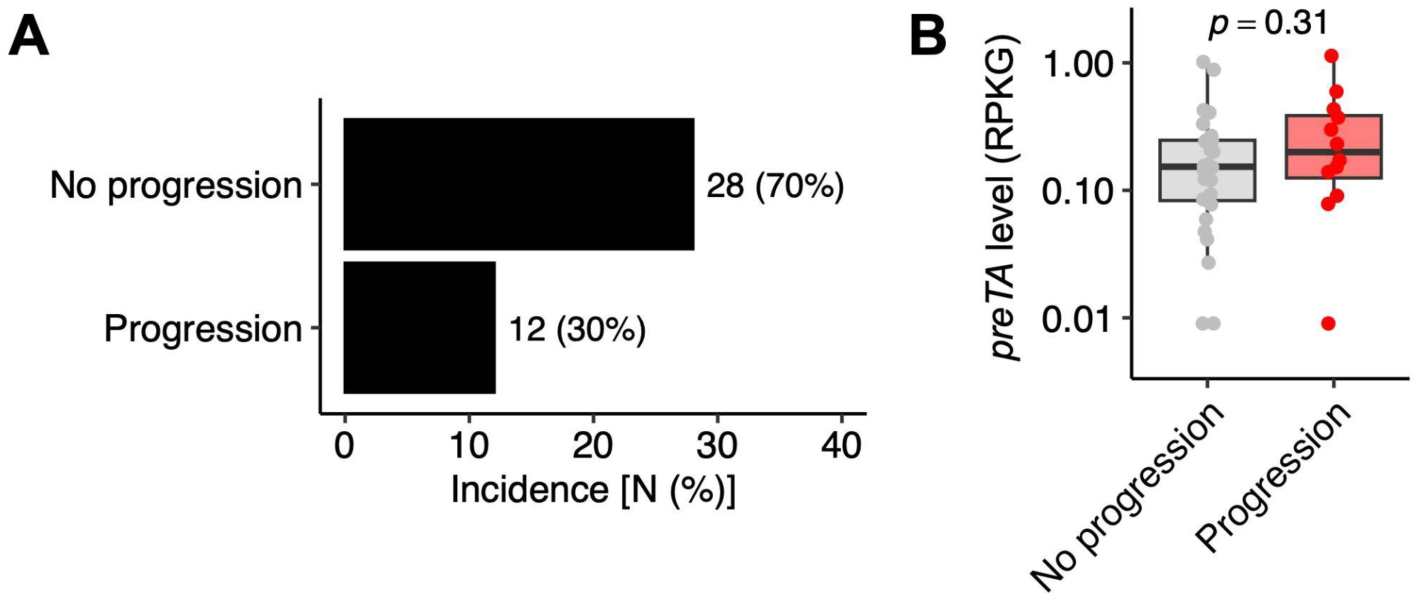
Microbial *preTA* is not associated with tumor progression. (**A**) Documented tumor progression during Cycles 1-3 in GO patients (*n* = 40). (**B**) Patients experiencing tumor progression have similar baseline stool bacterial *preTA* levels (*n* = 40). *p*-value: Mann-Whitney U test.

**Figure S17:**
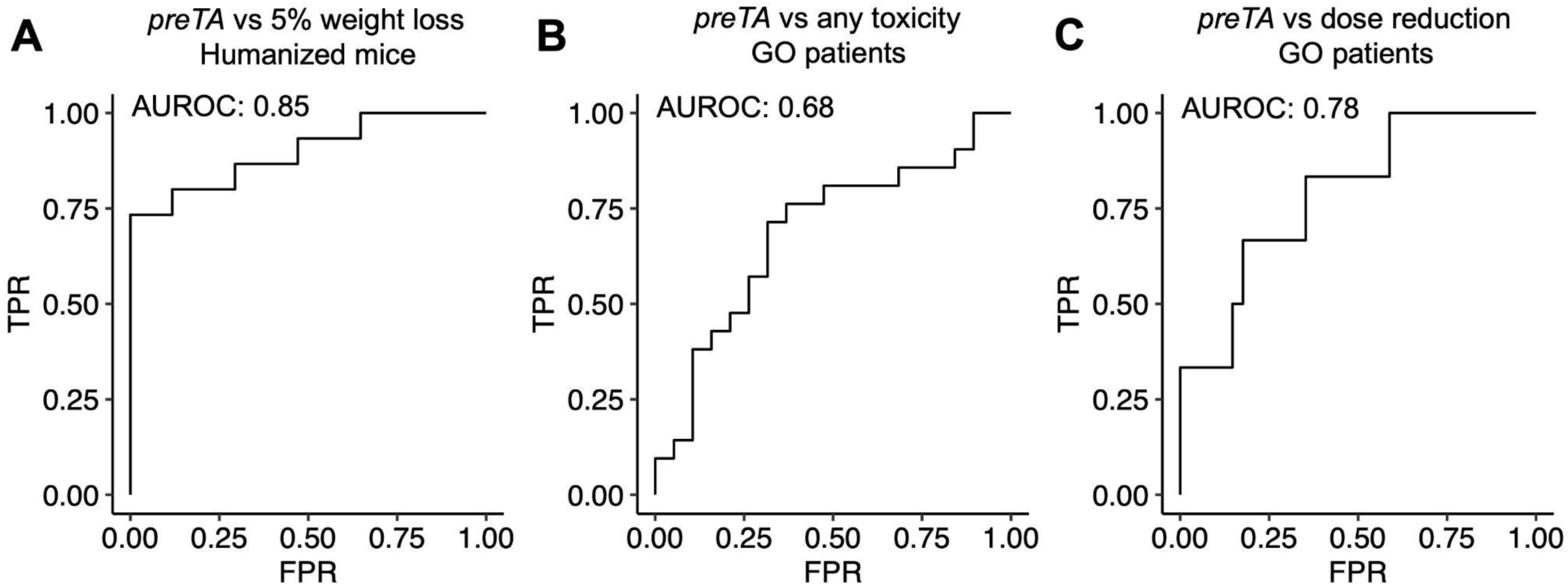
Microbial *preTA* predicts CAP toxicity. Univariate receiver operating characteristic (ROC) curves for classification of ≥ 5% weight loss in humanized mice (**A**), any documented toxicity in GO patients **(B)**, and dose reductions in GO patients **(C),** all using *preTA* level. True positive rate (TPR), false positive rate (FPR), and area under the receiver operating characteristic curve (AUROC) are reported.

**Table S1:** Baseline clinical characteristics of GO study patients. Values are displayed as median (IQR) for continuous variables and n for categorical variables. Data is for the 40 patients who submitted at least 1 stool sample.

|  |  |
| --- | --- |
| **Age (years)** | 52.5<br>(10.5) |
| **BMI (kg/m2)** | 24.8 (4.8) |
| **Sex** |  |
| Female | 20 |
| Male | 20 |
| **Race (self-reported)\*** |  |
| Asian | 4 |
| Black/African American | 2 |
| Unknown | 3 |
| White | 32 |
| **Ethnicity (self-reported)** |  |
| Hispanic or Latino | 3 |
| Not Hispanic or Latino | 34 |
| Unknown | 3 |
| **Cohort and concurrent pharmacologic treatment** |  |
| A: CAP as SOC | 21 |
| *CAP + oxaliplatin* | 9 |
| *CAP + radiation* | 9 |
| *CAP + bevacizumab* | 2 |
| *CAP + non-specified concurrent therapy* | 1 |
| B: TAS102 | 9 |
| *TAS102 monotherapy* | 5 |
| *TAS102 + bevacizumab* | 2 |
| *TAS102 + Y-90 radioembolization* | 2 |
| C: CAP + bevacizumab + pembrolizumab | 10 |
| *CAP + bevacizumab + pembrolizumab* | 10 |
| **Location of primary tumor** |  |
| Cecum/ascending | 7 |
| Transverse | 6 |
| Descending/Sigmoid | 11 |
| Rectosigmoid | 3 |
| Rectum | 13 |
| **TNM Stage at diagnosis** |  |
| I | 2 |
| II | 4 |
| III | 22 |
| IV | 11 |
| Unknown | 1 |
| **Prior surgical resection of primary** |  |
| Yes | 27 |
| No | 13 |
| **Ostomy** |  |
| Colostomy (present) | 2 |
| Colostomy (reversed) | 2 |
| Ileostomy (present) | 1 |
| None | 35 |
| **Metastases present** |  |
| Yes | 21 |
| No | 19 |

\*Total = 41 since 1 patient self-identified as multiple races.

**Table S2:** Mouse experiment metadata. Age is mean±sem at Day 0. CAP dose is daily total.

| Experiment | Group | n | Sex | Strain | Age (weeks) | Housing (n/cage) | LabDiet | CAP dose (mg/kg) |
| --- | --- | --- | --- | --- | --- | --- | --- | --- |
| Mechanisms of Toxicity | Vehicle | 8 | F | C57BL/6J | 9 | 4 | 5058 | 1500 |
| Mechanisms of Toxicity | CAP | 12 | F | C57BL/6J | 9 | 4 | 5058 | 0 |
| Antibiotic Depletion | Vehicle | 12 | F | C57BL/6J | 11.3 | 4 | 5058 | 1500 |
| Antibiotic Depletion | AVNM | 12 | F | C57BL/6J | 11.3 | 4 | 5058 | 1500 |
| Conventionalization | CONV-R | 8 | F | C57BL/6J | 8.1 | 4 | 5058 | 1100 |
| Conventionalization | GF | 8 | F | C57BL/6J | 13.6±0.3 | 4 | 5021 | 1100 |
| Conventionalization | CONV-D | 8 | F | C57BL/6J | 13.3±0.3 | 4 | 5021 | 1100 |
| CAP Regional Profiling Exp | Vehicle | 3 | F | C57BL/6J | 7 | 1 | 5058 | 0 |
| CAP Regional Profiling Exp | CAP | 3 | F | C57BL/6J | 7 | 1 | 5058 | 1000 |
| CAP Regional Profiling Exp | Vehicle | 2 | M | C57BL/6J | 7 | 1 | 5058 | 0 |
| CAP Regional Profiling Exp | CAP | 3 | M | C57BL/6J | 7 | 1 | 5058 | 1000 |
| SPF preTA Rescue Exp 2 | $\Delta preTA$ | 6 | F | C57BL/6J | 6.3 | 3 | 5058 | 1500 |
| SPF preTA Rescue Exp 2 | *preTA wt* | 6 | F | C57BL/6J | 6.3 | 3 | 5058 | 1500 |
| SPF preTA Rescue Exp 2 | *preTA*<sup>++</sup> | 6 | F | C57BL/6J | 6.3 | 3 | 5058 | 1500 |
| SPF preTA Rescue Exp 2 | $\Delta preTA$ | 6 | M | C57BL/6J | 6.3 | 3 | 5058 | 1500 |
| SPF preTA Rescue Exp 2 | *preTA wt* | 6 | M | C57BL/6J | 6.3 | 3 | 5058 | 1500 |
| SPF preTA Rescue Exp 2 | *preTA*<sup>++</sup> | 6 | M | C57BL/6J | 6.3 | 3 | 5058 | 1500 |
| SPF preTA Rescue Exp 1 | $\Delta preTA$ | 16 | F | C57BL/6J | 7.3 | 4 | 5058 | 1500 |
| SPF preTA Rescue Exp 1 | *preTA*<sup>++</sup> | 16 | F | C57BL/6J | 7.3 | 4 | 5058 | 1500 |
| SPF Stool PK | $\Delta preTA$ | 6 | F | C57BL/6J | 9.3 | 4 | 5058 | 1500 |
| SPF Stool PK | *preTA*<sup>++</sup> | 6 | F | C57BL/6J | 9.3 | 4 | 5058 | 1500 |
| SPF preTA qRT-PCR | $\Delta preTA$ | 4 | F | C57BL/6J | 9.3 | 4 | 5058 | 0 |
| SPF preTA qRT-PCR | *preTA wt* | 4 | F | C57BL/6J | 9.3 | 4 | 5058 | 0 |
| SPF preTA qRT-PCR | *preTA*<sup>++</sup> | 4 | F | C57BL/6J | 9.3 | 4 | 5058 | 0 |
| Gnoto preTA Rescue | GF | 6 | M | C57BL/6J | 7.5±0.4 | 1-3 | 5021 | 1500 |
| Gnoto preTA Rescue | $\Delta preTA$ | 6 | M | C57BL/6J | 8.1±0.4 | 1-3 | 5021 | 1500 |
| Gnoto preTA Rescue | *preTA*<sup>++</sup> | 6 | M | C57BL/6J | 8.1 | 1-3 | 5021 | 1500 |
| GO Humanization Exp 1 | Donor 35 | 8 | M | C57BL/6J | 9.8±0.5 | 1-3 | 5021 | 1500 |
| GO Humanization Exp 1 | Donor 32 | 8 | M | C57BL/6J | 9.7±0.5 | 1-3 | 5021 | 1500 |
| GO Humanization Exp 2 | Donor 7 | 8 | M | C57BL/6J | 11.5±0.1 | 1-3 | 5021 | 1500 |
| GO Humanization Exp 2 | Donor 39 | 8 | M | C57BL/6J | 11.4±0.1 | 1-3 | 5021 | 1500 |

**Table S3:**
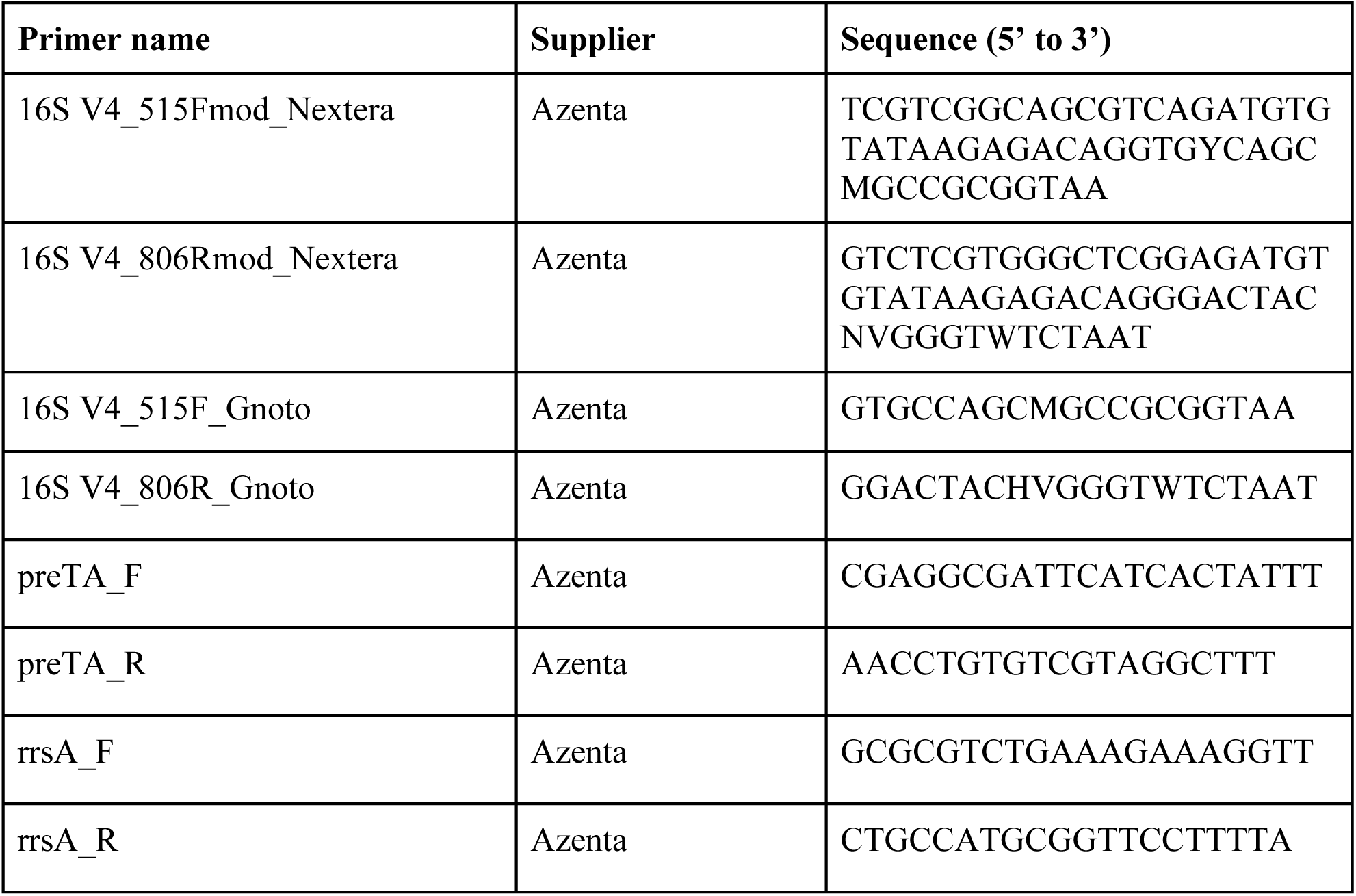
Primer sequences.

## Notes

### Summary of Updates

This version of the manuscript has been revised to include additional experiments studying the effect of CAP along the gastrointestinal tract in mice, the effect of non-engineered bacterial strains on CAP toxicity, conservation of CAP-microbiome interactions across sexes, and pharmacokinetic experiments demonstrating that preTA++ E. coli lowers detectable 5-FU in stool.

